# Adaptive rewiring and temperature tolerance shape the architecture of plant-pollinator networks globally

**DOI:** 10.1101/2025.10.19.683289

**Authors:** Gaurav Baruah, Meike J. Wittmann

**Author notes:** Faculty of Biology, Theoretical Biology, University of Bielefeld, 33501 Bielefeld, Germany.

## Abstract

Rising environmental temperatures are rapidly reshaping plant–pollinator communities by altering species traits and interaction patterns. We develop a simple eco-evolutionary model that integrates species-specific temperature tolerance curves with phenotype-based interaction dynamics. Across temperature gradients, species adaptively rewire, that is, they change their interaction partners. This rewiring is an emergent property of our model, driven by temperature-mediated selection and co-evolutionary trait matching. As temperature increases, our model predicts a consistent decline in network-level specialization, alongside increasing connectance and nestedness which are signatures of structural re-organization. These predictions are supported by empirical patterns from 165 plant–pollinator networks worldwide, where mean annual temperature correlates positively with connectance and nestedness, and negatively with network specialisation. Our findings suggest that temperature-driven trait evolution and emergent adaptive rewiring govern the assembly and architecture of mutualistic networks. By bridging dynamical eco-evolutionary theory with global empirical data, this work reveals the central role of trait-based processes in structuring biodiversity under ongoing and accelerating climate warming.

## 1 Introduction

One of the fundamental goals of community ecology has been to understand the causes and consequences of species interactions for the maintenance of biodiversity and ecosystem functioning across space and time ^1–3^. Traditionally, species interaction networks have been treated as static and this static view has no doubt advanced our understanding of how communities function and maintain diversity ^4–10^. However, it is now clear that species create and sever links dynamically in response to changing densities, phenotypes, and environmental conditions ^11–15^. For example, after the loss of a resource or a partner species due to changes in the environment ^16,17^, species sometimes shift to another interaction partner through shifts in traits. This shift in species interaction i.e., “interaction rewiring” ^8,13,18^ could thereby impact the stability of the community.

In plant-pollinator mutualistic systems, for example, trait and phenological matching is essential for effective pollination. ^19,20^. Seasonal shifts in temperature or precipitation can rapidly reconfigure who interacts with whom ^13,21–23^. Recent empirical and theoretical work has documented “rewiring” of interactions over time ^13,17,22,24–26^, yet we still lack a clear mechanistic understanding of how these dynamic patterns propagate through ecological networks and influence biodiversity patterns and responses to environmental change ^13^. Unpacking these drivers is critical if we are to predict and manage the resilience of ecological communities in an era of rapid global change.

Temperature is a fundamental environmental variable that varies predictably along latitudinal gradients and shapes global biodiversity patterns^27–29^. Near the equator, temperature is relatively stable throughout the year, while at higher latitudes, organisms are exposed to greater seasonal fluctuations. This gradient influences the evolution of species’ thermal tolerances, with low-latitude species often having narrower thermal windows than those at temperate or polar regions ^27–30^. Such variation in temperature tolerances can strongly affect how species respond to climate change. In plant–pollinator communities, temperature not only governs species’ survival and activity periods but also shapes the synchrony and stability of species interactions ^31–33^. These communities are especially vulnerable to temperature changes because both partner species i.e., plants and pollinators, must be active concurrently for successful pollination to occur. Insect emergence, flowering time, and interaction strength are all tightly linked to temperature, and mismatches in these events can disrupt network integrity^33–35^.

Plant-pollinator communities exhibit a characteristic structure quantified by metrics such as connectance, which measures the proportion of realised links out of all possible links. Previous research has consistently demonstrated the existence of pronounced latitudinal gradients in the organization and structure of ecological interaction networks^36–38^. Despite these advances, a critical knowledge gap remains: the precise ecological and evolutionary mechanisms that govern and shape the organization of such complex networks across spatial and environmental gradients are still poorly understood. Such characteristic architecture of plant-pollinator networks could potentially emerge from a combination of ecological and evolutionary constraints driven by response to abiotic environmental conditions (Fig. 1)^39–41^. Since temperature tolerance could potentially shape the temporal activity, it could directly influence who interacts with whom, thereby influencing species rewiring and shaping the architecture of such communities (Fig.1). Modeling such temperature-driven dynamics is thus critical for predicting how warming may alter not only species distributions but also the architecture and stability of ecological networks ^42^. Gaining a foundational understanding of how species interaction dynamics influence the temporal dynamics and structural organization of networks, especially in relation to changing environmental conditions, is essential for uncovering how complex communities assemble and persist over time. Such insights are crucial not only for predicting the response of individual species but also for anticipating how entire networks may reorganize under environmental change.

**Figure 1:**
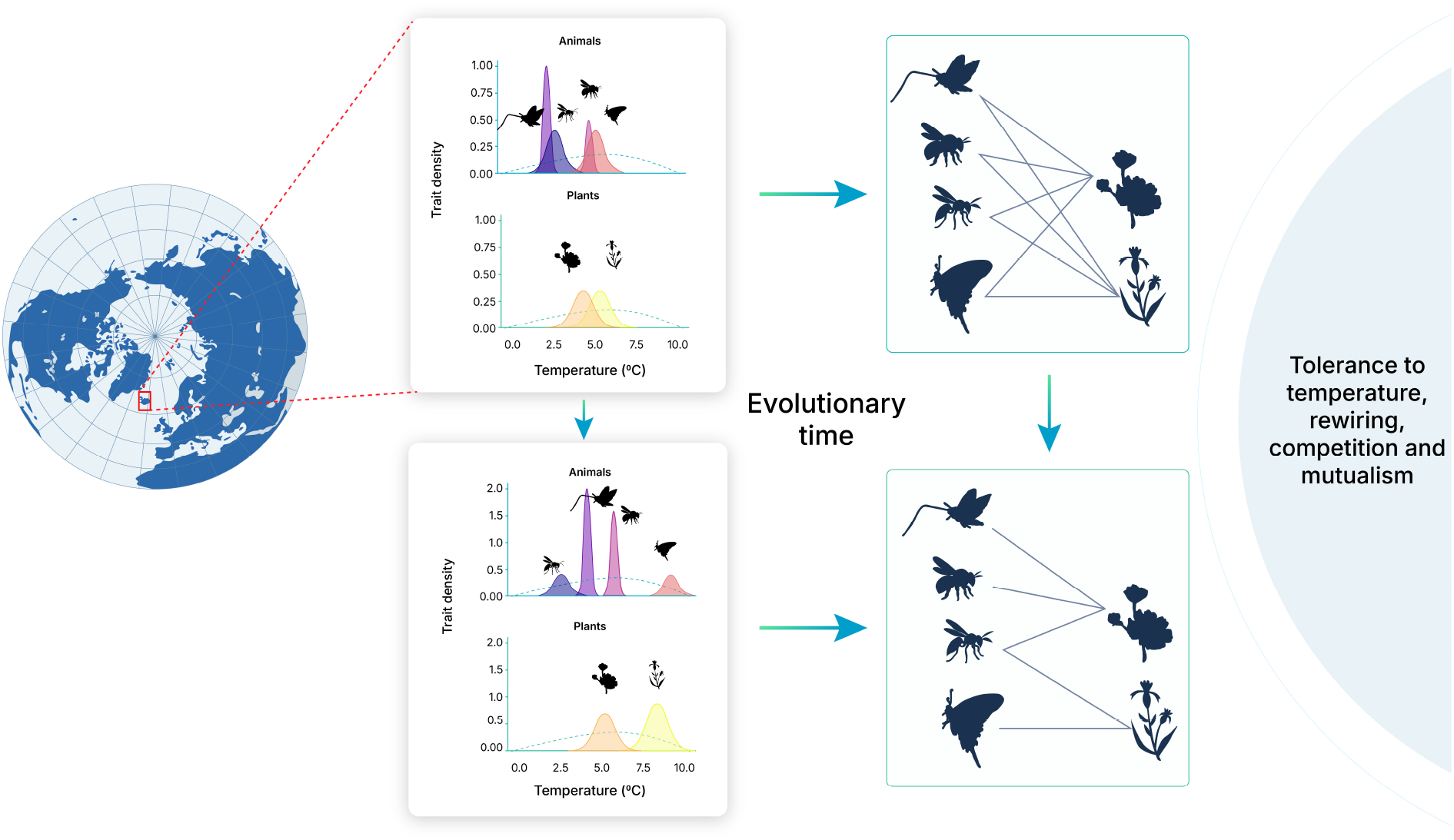
Graphical illustration of how biotic and abiotic conditions shape the architecture of a plant-pollinator network. (First row) A hypothetical plant-pollinator network of six species found in a subpolar region. The resulting six species initially could have similar optimum phenotypes such that all pollinators can potentially interact with all plants as depicted by the structure of the network. The dashed curve depicts the hypothetical temperature-tolerance curve under which species in the sub-polar region could have positive growth. (Bottom row): Over eco-evolutionary time, species adaptively rewire in the phenotype axis in a way to have minimal competition and higher mutualistic trait overlap dictated by environmental conditions such as temperature, which re-structures the species network.

In this study, we develop a simple model of eco-evolutionary feedbacks in plant-pollinator networks, driven by phenotype-based processes and species’ tolerance to temperature. Adaptive species rewiring emerges from this framework and shapes the architecture of such networks across a temperature regime. Our model integrates quantitative genetics and trait evolution to explain how competition and mutualism shape interactions across temperature gradients. It predicts a decline in network specialization with rising temperature, consistent with global latitudinal patterns of plant–pollinator networks. In addition, plant-pollinator networks become more nested and more connected as temperature increases. Our model was able to successfully reproduce the specialization and architectural patterns observed in 165 empirical plant–pollinator networks worldwide across a latitudinal and temperature gradient. Together, our results demonstrate that species’ thermal-tolerance, co-evolutionary trait matching, and emergence of adaptive rewiring jointly play a major role in how plant-pollinator networks structure and assemble over time.

## 2 Results

In our eco-evolutionary model (see Methods, equation 1, 5, 6), phenotype-based competitive and mutualistic interactions, along with selection pressures driven by temperature, influence the resulting network architecture of plant-pollinator interactions (see conceptual Fig. 1). We modelled optimum phenotype as a continuous trait determining both mutualistic interactions and competitive overlap. Individuals with similar optimum phenotypes could interact positively (plant–pollinator) or compete (pollinator–pollinator). We simulated plant-pollinator interactions across a temperature gradient ranging from 0°C to 29°C, running eco-evolutionary simulations for (10^4^)time units for each temperature (see Methods). These different temperature regimes mimicked MAT (mean annual temperature) across the latitudinal gradient of the compiled empirical plant-pollinator networks.

### 2.1 Plant-pollinator eco-evolutionary and interaction dynamics in response to temperature

In our simulations, all plant and pollinator species theoretically can interact, so baseline interaction probabilities are non-zero. Species then sort themselves according to how their optimal phenotypic values align with their partner species, under the impact of resource competition, mutualistic interaction, and tolerance to environmental temperature (see a graphical description in Fig. 1). In Fig. 2, the network of twenty-two plants and pollinators, initial connectance of the artificial network are based on how much their optimum phenotypes overlap based on a trait-similarity matrix (see Methods section 5.1 for quantification of the trait-similarity matrix). Initial connectance of the twenty-two species artificial plant-pollinator in the upper row network was 0.14.

**Figure 2:**
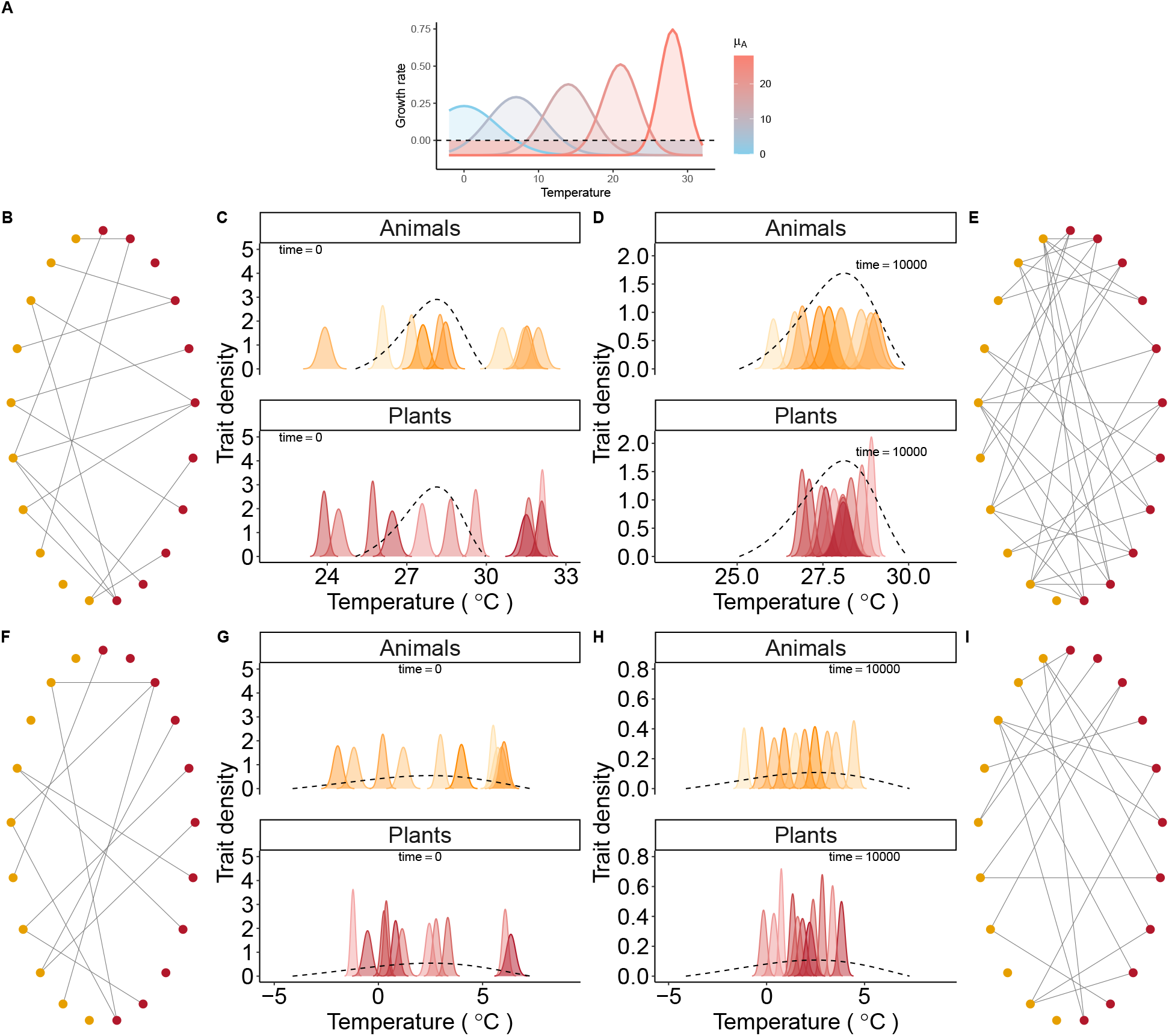
Adaptive phenotypic rewiring in response to temperature organises the architecture of a plant-pollinator network. A) Different growth rates of species at different temperature given by *a* =0.1 here; *a* controls the shape of the temperature tolerance curve, with *a >* 0 indicating that tolerance to environmental temperature is dependent on species optimum trait. Different colors show species with different mean optima. Blue color indicates species with a low temperature optimum akin to a species that might occur at higher latitudes i.e., at sub-polar/arctic regions for instance. (B) A network of 22 species with starting network connectance of 0.14 based on initial trait overlap. Red nodes represent plants and orange nodes represent pollinators. C) Starting trait distribution of plants and pollinators and D) final trait distribution of plants and animals at quasi eco-evolutionary equilibrium. E) Resulting plant-pollinator network connectance increases to 0.35 from 0.14 for an environmental temperature at 28° C. (F-G-H) Another network with the same number of species with initial connectance of 0.15. At eco-evolutionary equilibrium, connectance slightly increases to 0.175 for an environmental temperature of 2° C. The dashed curves in C, D, G, and H show the average per-capita temperature tolerance curve for all species, calculated from Equation 4 at 28° C or 2° C, where *µ* represents the mean optimal trait value across species in the network.

Tolerance to environmental temperature was species-specific in our model (Fig. 2A). In our model, temperature tolerance shape was modulated by a parameter *a* and the species’ average optimum trait value (see figure S1). When *a >* 0, a higher average optimum phenotype of a species (e.g. a species adapted to tropical temperatures) leads to a narrower thermal tolerance curve. For instance, in our model, a network of twenty-two species found in tropical latitudes implies that these species have high optimum trait values. Consequently, their temperature tolerance curves are narrow (*a >* 0; Fig. 2B–E), reflecting the low environmental variability typical of tropical regions ^28^. In contrast, species in temperate or polar latitudes exhibit lower optimum trait values and broader tolerance curves (*a >* 0), consistent with higher environmental variability (Fig. 2F–I). However, when *a* =0, the tolerance curve maintains a constant shape across latitudes, whereas when *a >* 0, both the peak and breadth of the curve vary with temperature—narrowing in warmer regions and widening in cooler ones (Fig. S1)

We observed that at eco-evolutionary equilibrium, network connectance increased for such a twenty-two species network under high temperatures increased to 0.35 from 0.14, such as analogous to those observed in tropical latitudes with little environmental variability (small thermal tolerance widths, *a >* 0) (Fig. 2B–E); but connectance of a similar twenty-two species network declines under cooler, higher-latitude environmental temperatures (Fig. 2F–I), such as those observed in temperate to arctic/polar regions (high thermal width, and low thermal peak, *a >* 0). At eco-evolutionary equilibrium in Fig. 2, network connectance remains around 0.17 from 0.15 given by the optimum-trait similarity matrix estimated based on Gower-variance similarity matrix (see methods).

### 2.2 Adaptive species rewiring to environmental temperature

Within our modeling framework, species were capable of adaptively switching their interaction partners i.e., rewire. Unlike previous models that imposed rewiring whenever interactions became beneficial^18,26^, here partner switching emerges naturally from the joint evolution of mean phenotypes under temperature-driven selection and competition pressures (Fig. 1). We quantify rewiring by comparing successive community “snapshots”, calculating phenotype-based similarity matrices of species between each plant’s trait distribution and each pollinator’s, and recording a new link whenever similarity as well as species densities surpass predefined thresholds (see Methods for details).

We found that adaptive phenotypic rewiring increased over eco-evolutionary time, but was modulated by both environmental temperature and competitive strength among species (Fig. 3A). When environmental temperature increased, the median magnitude of adaptive rewiring increased, particularly in networks characterized by *a >* 0, i.e.,when the shape of the temperature tolerance curve was dependent on the mean optimum of a species (Fig. 3B). However, this also depended on the starting conditions (Fig. S12), supposedly because less rewiring is needed if the spread of the starting mean trait values is already similar to the equilibrium for the given temperature. These results show that when adaptive rewiring couples with temperature filtering, it dynamically restructures plant-pollinator networks networks.

**Figure 3:**
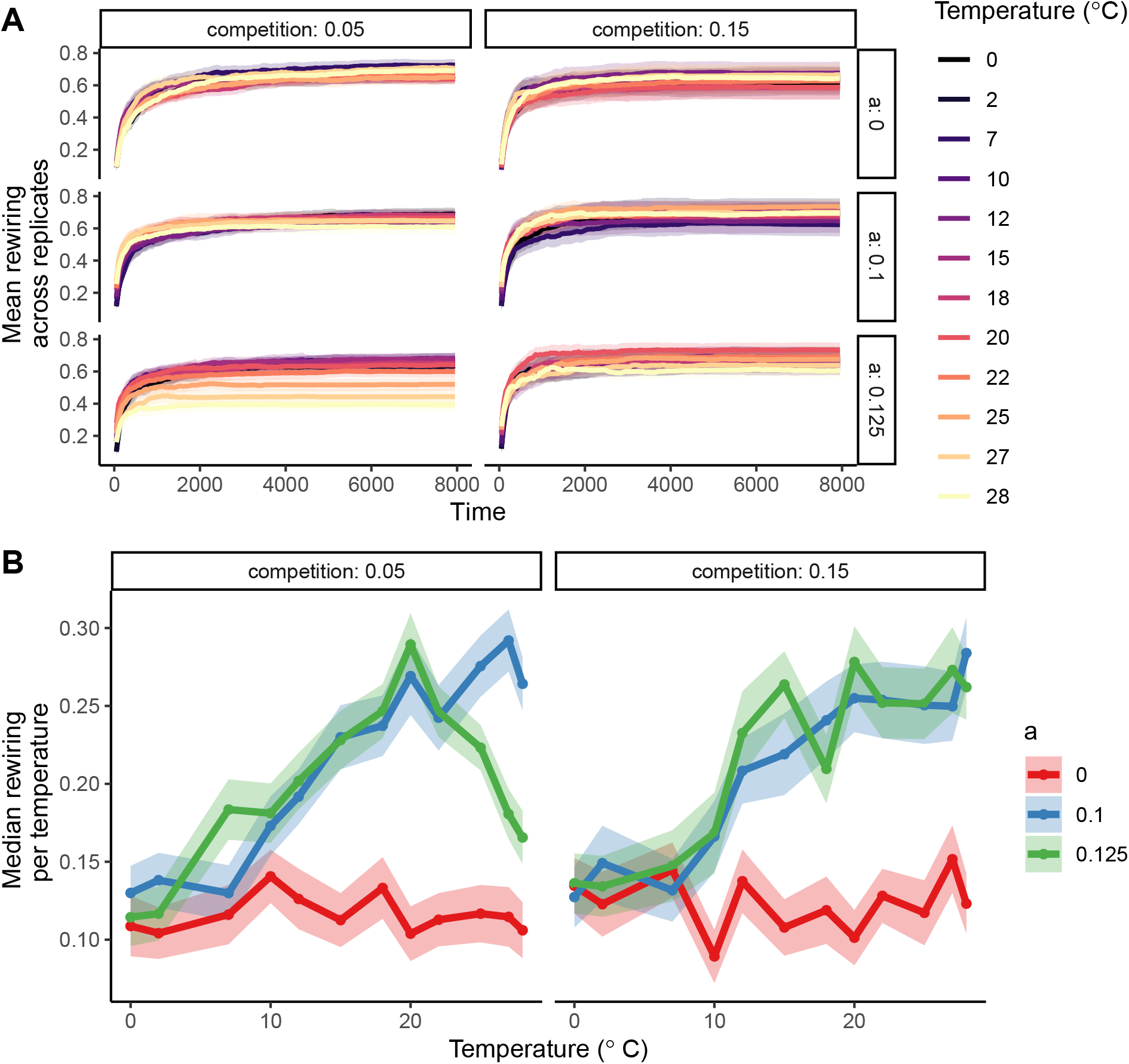
Dynamics of adaptive phenotypic species rewiring. A) Mean rewiring over time across replicates for a simulated plant-pollinator network with 20 species (shown only for 8000 time units). B) Median phenotypic rewiring in relation to temperature, competition, and shape of the temperature tolerance curve given by *a*. Median rewiring increases as temperature increases only when *a >* 0. On average, with increasing competition strength, median rewiring slightly increased. Lines represent median rewiring per environmental temperature and shaded areas represent standard error across replicates. Rewiring was estimated based on the trait overlap matrix given by the Gower variance metric (see Methods and Models). Parameters as in Table S1.

### 2.3 Network architecture in response to environmental temperature regimes

When *a* =0, i.e., the shape of the temperature tolerance curve is independent of species mean optimum trait, equilibrium connectance, nestedness (NODF), modularity, and network specialisation (H2’) had no relationship with environmental temperature (Fig. 4A-D, *a* =0). However, when *a >* 0, i.e.,when the shape of the temperature tolerance curve was dependent on the mean trait of species, equilibrium network architecture was impacted by environmental temperature. In particular, at eco-evolutionary equilibrium, connectance and nestedness were positively associated with environmental temperature (Fig. 4A-B, *a* =0.1, *a* =0.125). Equilibrium network modularity and network specialisation (H2’) were negatively related to temperature (Fig. 4C-D, *a >* 0). These results were robust to the choice of similarity metrics used to characterise and quantify network architecture (see Fig. S5-S10), different thresholds for interaction (see Fig. S15-S16), or different plant-pollinator network sizes (see Fig. S7-S10). Strength in competition dictated by the competition width impacted the overall intercept rather than the slope of the relationship in all the cases. Moreover, our modelling framework implicitly tested latitude-linked diversity gradients^37^ by assessing how plant richness shapes plant–pollinator network architecture (see Methods and appendix S2). Although we modelled elevated plant richness in tropical temperatures, we nonetheless detected a negative association between mean annual temperature and network specialization H2’ (Fig. S4).

**Figure 4:**
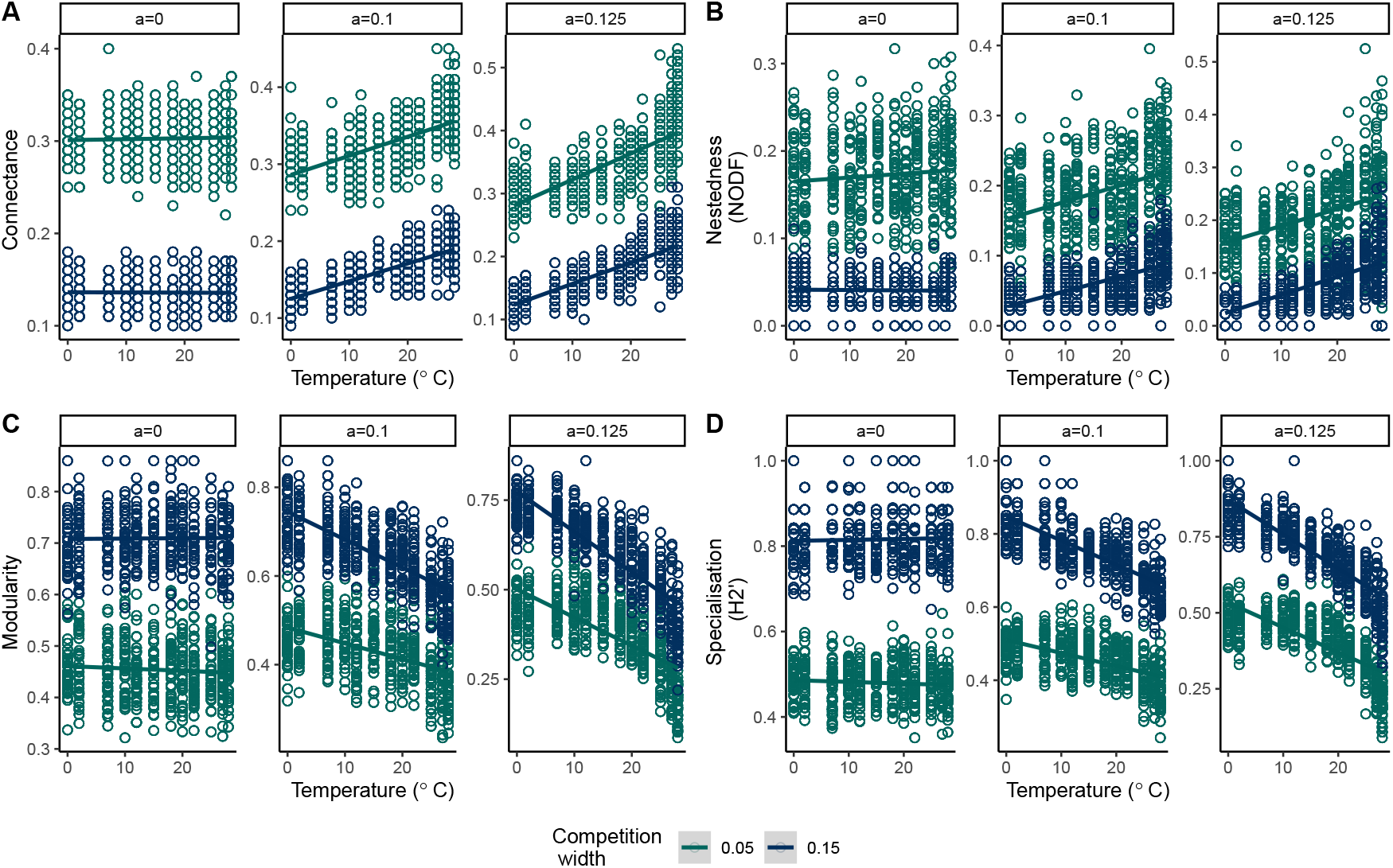
Adaptive phenotypic species rewiring and tolerance to temperature impact network specialisation and structure of plant-pollinator networks. A) As temperature increases, final equilibrium network connectance increases when *a >* 0, with strong competition causing an overall decrease in network connectance. B) Similarly, nestedness increases as temperature increases when *a >* 0. C) Modularity was negatively correlated with temperature when *a >* 0. D) H2’, network specialisation, decreases as temperature increases when *a >* 0, with competition having an impact more on the intercept and less on the slope of the relationship. Lines represent linear model fit. All metrics are based on networks determined from the modified Gower variance metric. Starting network size is 20. Parameters as in Table S1.

### 2.4 Architecture of empirical plant-pollinator networks across a climatic gradient

We compiled 165 plant-pollinator network data globally (Fig. 5A) along with mean annual temperature (MAT) data at the network locations (see Methods section 4.4). Principal component analysis (PCA) of the network metrics such as connectance, nestedness (NODF), modularity revealed that the first principal component axis was positively correlated with nestedness (Pearson correlation coefficient, 0.90), and connectance (Pearson coefficient 0.93), and negatively to modularity (pearson coefficient =−0.85) and network size (Pearson coefficient of −0.78) (Fig. S2). Since these network metrics were highly correlated, we used PC1 axis scores as predictor of network architecture.

**Figure 5:**
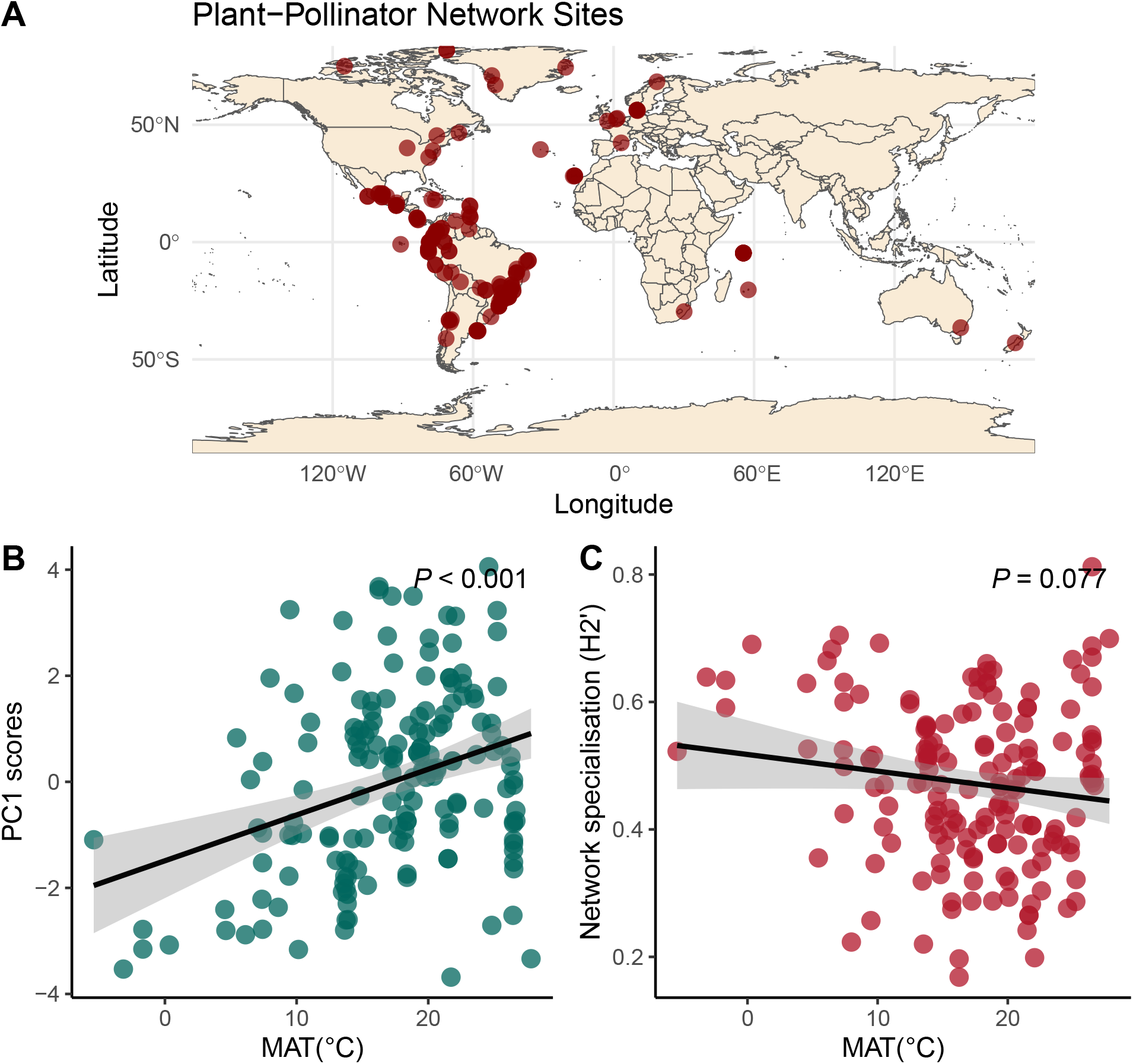
Network architecture of plant-pollinator networks in relation to mean annual temperature. A) Geographic location (latitude and longitude) of 165 plant–pollinator networks. (B) Relationship between principal component axis 1 (PC1) scores and mean annual temperature (MAT). PC1 scores were positively correlated with nestedness and connectance, and negatively correlated with network size and modularity. (C) Network specialization (H2) in relation to MAT. Each point represents a plant–pollinator network. Lines indicate fitted ordinary linear regressions with 95% confidence intervals.

Similar to our modelling results, we observed a strong positive relationship between mean annual temperature (MAT) and PC1 scores (*R*^2^ =0.19, *p <* 0.001, Fig. 5B), indicating that network connectance and network nestedness increased as mean annual temperature increased, and modularity and network size decreased as mean annual temperature increased (Fig. 5B). Furthermore, we also observed a negative but nonsignificant relationship between network specialisation (H2’) and mean annual temperature (Fig. 5C), consistent with our modelling results. In addition, we observed similar associations between plant-pollinator network metrics and mean annual temperature after removing spatial correlation among network site locations using a spatial eigenvector filtering approach (see Methods, and appendix S3, and figure S17).

In addition, our model produced a negative relationship between network size (i.e., total species in a network) and final network connectance at eco-evolutionary equilibrium (Fig. S11), modulated by the the shape of the temperature tolerance curve (i.e., *a*). This negative relationship between network size and connectance was also observed empirically in the dataset of 165 plant-pollinator networks (Fig. S2A PCA biplot), and is an emergent property that has been widely observed in other datasets of plant-pollinator networks ^43^.

## 3 Discussion

Increasing environmental temperatures threaten biodiversity and compromise the stability of complex ecosystems ^32,44^, with plant–pollinator networks among the most temperature-sensitive systems ^33,45^. Shifts in ambient temperature can disrupt plant–pollinator interactions and impact community resilience ^33,34,45,46^. In this study, we develop an eco-evolutionary dynamical model based on quantitative genetics ^41,42^, and evaluated how environmental temperature and phenotype-based co-evolution^13,14,17,25^ impacted the structure and assembly of plant-pollinator networks. In addition, using 165 plant-pollinator networks distributed globally, we show that our modelling results qualitatively predicted global empirical patterns of network architecture. Our model is set up to capture many crucial factors that shape the architecture of plant-pollinator networks. In our models, all plants and pollinators could interact with one another, but individuals with similar optimum phenotypes had stronger mutualistic benefits^47,48^. This reflects empirical evidence that interactions are more likely when species are active under similar environmental conditions, such as temperature-driven phenological overlap^33,49–52^. We modelled optimum phenotype as a continuous trait determining both mutualistic interactions and competitive overlap. Individuals with similar optimum phenotypes could interact positively (plant–pollinator) or compete (pollinator–pollinator). All species started with random positions on the optimum phenotype axis. Over eco-evolutionary time, trait-based interactions and temperature-dependent processes drove species to self-organize along the trait axis, reorganizing plant-pollinator architecture. Thus, adaptive phenotypic rewiring was an emergent phenomenon. This could occur when such species face strong within-guild competition and/or face narrow environmental niches caused by the environmental temperature^28,29,53^.

As the optimum phenotype increases, a species’ thermal-tolerance curve sharpens, parameterized by *a*. If *a* =0, the temperature tolerance curve for a species with a higher optimum phenotype has the same shape as for a species with a lower optimum phenotype. With *a >* 0, the temperature tolerance curves has increasingly narrow widths with increasing species optimum phenotypic value, similar to what was observed empirically^27,28^. The narrow thermal niches at elevated temperatures, typical of tropical systems, were characterised by the thermal tolerance curves with *a >* 0 and high species mean optimum trait value^28,53^. When *a >* 0 and when there was relatively high among-species variation in mean optimum traits, rising environmental temperatures triggered marked increases in species rewiring, driving up connectance and nestedness while reducing specialization and modularity. Such narrow tolerance curve for species having higher optimum phenotypes, akin to those found in tropical climates, forces species phenotypic optima to cluster, resulting in greater niche overlap. Consequently, pollinators with high thermal optimum can exploit different niches of plants at those temperatures, lowering specialization but boosting connectance and nestedness. However, intense interspecific trait-based competition (competition width =0.15) could constrain such plant-pollinator overlap, enforcing exclusive mutualistic interactions, which heightens specialization and decreases both connectance and nestedness (Fig. 4A-B,4D) while still maintaining a positive relationship between network structure (connectance, nestedness) and temperature.

At tropical latitudes, where environmental temperatures fluctuate less than in temperate and arctic zones ^28,53,54^, species experience relatively stable conditions that reduce selective pressure for broad tolerance widths, in contrast to the greater thermal variability at higher latitudes that favors wider tolerance ranges. Previous macroecological analysis of similar mutualistic plant-pollinator and seed-dispersal assemblages showed that networks along the tropics become more nested, similar to what we observed both in our theoretical and empirical results^36–38^.Our modelling results, and analysis of empirical network data, demonstrate that plant-pollination assemblages exhibit positive correlation between mean annual temperature on the one side and connectance or nestedness on the other side. In bipartite mutualistic networks, nestedness reflects a core–periphery organization in which generalists interact with many partners, and specialists engage only with proper subsets of those generalists’ partners^7,55–57^. Connectance, which is the proportion of possible links that are realized, fuels this structure: as overall connectance rises, generalists accumulate additional interactions, enlarging the link pool from which specialists interact and thereby accentuating the nested pattern. Under elevated thermal regimes (e.g. tropical conditions) and weak competition, species’ thermal-tolerance curves constrict, promoting trait convergence and greater niche overlap. This drives up connectance, and in turn nestedness increases as these generalists form a robust network core with specialists, by virtue of their subset interactions.

We also observed a negative correlation between mean annual temperature and network specialization, both in the theoretical models and in the empirical dataset of 165 networks (though not significant there). This negative relationship in network specialization and MAT has been previously attributed to higher plant species diversity in the tropics, which lowers per-species resource abundance and thus promotes generalist behavior, while temperate or arctic communities with fewer resource plants enforce narrower interaction niches ^37^. Here, we tested this mechanism by simulating communities in which pollinator richness remained constant but plant richness declined toward higher latitudes or colder climates, and measured specialization under low-temperature to high-temperature regimes (see supplementary appendix S2 and figure S4). If species diversity drove network generalization at warmer, low-latitude sites, then network specialization would decline with increasing temperature even when *a* =0 (no change in thermal despite warm or cold temperatures). Instead, in Figure S4A, specialization actually rises with temperature for *a* =0, and connectance correspondingly decreases (Figure S4B), regardless of competition strength. In contrast, when *a >* 0, i.e., with species-specific thermal tolerance curve, network specialization decreases as temperature increases despite higher plant richness, consistent with the empirical observations. This suggests that the shape of thermal tolerance curves, together with competition and mutualistic interactions, was the primary mechanism driving generalized interactions under warm temperature regimes.

We theoretically show that temperature-induced narrowing of thermal-tolerance niches that drives phenotypic shifts and adaptive rewiring ^13,58^ can structure plant-pollinator networks across a temperature gradient. Thermal tolerance niches and rewiring produces comparable increases in network generalization even when species richness is held constant. Because our approach controls for initial network size, observed differences in specialization and network architecture in figure 4 arise solely from phenotypic driven shifts governed by the shape of the thermal tolerance curve, competition, and mutualistic trait-matching co-evolution. Specifically, we found that when *a >* 0, adaptive rewiring emerges to impact species phenotypic overlap and drives changes in specialization independently of species richness. These results show that temperature-mediated changes in niche width and adaptive rewiring can reproduce latitudinal patterns of network specialization without invoking differences in diversity. These theoretical results match empirical observations across multiple globally mutualistic communities.

We do, however, acknowledge that there are several aspects that we could not theoretically test, one of which is the impact of precipitation, species dispersal, or human impacts ^59^. One interesting avenue for future research would be to theoretically test the impact of anthropogenic pressures by inducing an external mortality in such models and link it with MAT or with absolute latitude values. Species dispersal is another factor that we ignored, which has been suggested to be an important contributing factor in maintaining biodiversity and structure of ecological communities ^60,61^. In addition, we also acknowledge that our model was unable to test effects of seasonality of temperature on such interactions^13^. We do encourage future studies to explicitly look at how these factors shape the architecture of plant-pollinator networks.

## 4 Methods and Models

### 4.1 Modelling framework: Plant-pollinator mutualistic dynamics

We model the dynamics of pollinators and plants in a quantitative trait *z* that relates to its optimum temperature (we refer to it as optimum phenotype). Within each plant or pollinator species *i*, the trait is normally distributed with density function 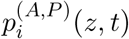, with mean *µ*_*i*_ and phenotypic variance 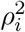, and with each individual having its own trait value *z*. Here the superscripts *A* and *P* stand for pollinators and plants, respectively. The governing dynamical equations of population dynamics can be written with modified Lotka-Volterra equations. The per-capita growth rate of pollinators, 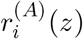, and plants, 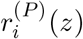, can be written as modified from ^42,62,63^ (see details in supplementary appendix)

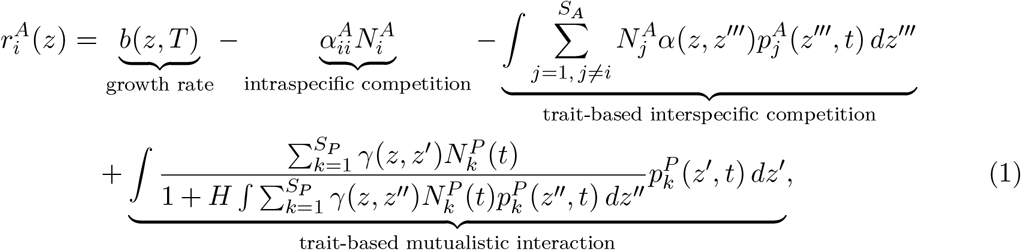

where 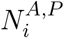 represents the population density of species *i* in the animal or plant guild, respectively. The term *b*(*z, T*)denotes the temperature-dependent growth rate of an individual with phenotype *z*, independent of competitive or mutualistic interactions. This formulation assumes that both plants and pollinators are similarly affected by local temperature *T*. The parameters *S*_*A*_ and *S*_*P*_ represent the total number of pollinator and plant species, respectively. *H* is the handling time parameter.

The term 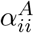 corresponds to the strength of intra-specific density dependence for species *i* within the pollinator guild (and analogously for plants). This terms captures all the other aspects of species growth rate that might not be captured by the individual’s optimum trait. Based on previous studies, this self-regulation parameter is an important aspect in determining the stability of species interaction networks^64^. To that end, we fixed intra-specific competition *α*_*ii*_ to be at 1, which in our framework was greater than inter-specific competition (see table of parameters S1). On the other hand, *α*(*z, z*_*′′′*_)captures the intensity of trait-based inter-specific competition between individuals of different species within the same guild (either plants or pollinators). This is modeled using a Gaussian competition kernel:

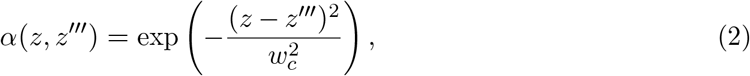

implying that pollinators with similar optimal temperatures — and thus overlapping phenologies — experience stronger competition for floral resources blooming at those times.

The mutualistic interaction between individuals is captured by the kernel *γ*(*z, z*^*′*^), which quantifies the interaction strength between a pollinator individual with thermal optimum *z* and a plant individual with thermal optimum *z*^*′*^. This interaction can also be modeled as a Gaussian function:

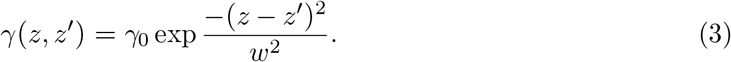

In this equation, *γ*^0^ represents the maximum strength of mutualistic interactions, while *w* controls the width of interaction between individual phenotypes. When the optimal temperatures of a plant and a pollinator are similar, mutualistic benefits are enhanced. Biologically this means that the flowering temperature of plants matches the emergence temperature of pollinators, increasing the likelihood of mutualistic interaction^40,41,47^. Temperature strongly influences species phenology, such as plant flowering times and pollinator emergence, which are often tied to temperatures. Thus, mutualistic interactions are more likely when their optimal temperature traits align ^33^.

The temperature-dependent growth or tolerance curve is a modified version of the curves in studies^41,65^ but see ^66^:

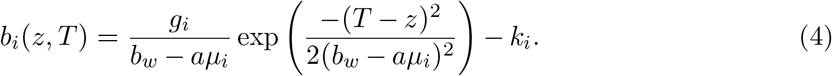

In this equation, *T* represents the local temperature, *µ*_*i*_ is the mean optimum temperature of species *i*, while *b*_*w*_, *g*_*i*_, and *a* are parameters that shape the temperature tolerance curve (see figure S1); *k*_*i*_ is the intrinsic mortality rate, which was fixed at 0.1. The parameter *b*_*w*_ determines the width of the curve, and the ratio 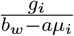 influences the height of the peak of the curve^65,66^, and *k*_*i*_ is the intrinsic mortality. If *a >* 0, the shape of the tolerance curve is also impacted by the mean optimum trait of the species. If the mean optimum of the species is low, the shape of the tolerance curve is wider and the fall-off from the optimum temperature is slower. However, if the mean optimum phenotypic trait *µ*_*i*_ of a species is high, the shape of the tolerance curve becomes narrower, and the fall off from the mean is sharper (see figure S1). This specific formulation follows from empirical studies where it has been consistently shown that there is a correlation between tolerance width and latitude i.e., at higher latitudes there is higher temperature variability than at lower latitudes, where temperature is less variable. As a consequence, species at higher latitudes tend to have a broader thermal tolerance width in contrast to species at lower latitudes^28^. We, however, do not model temperature variability. Instead, we explore this scenario by introducing (*b*_*w*_ − *aµ*_*i*_)which then mimics the fact that populations at higher latitudes, which are exposed to more variable temperature fluctuations, are adapted to such variability by having a wider temperature tolerance width. This type of formulation closely associates with growth curves found empirically^27,28,30,53,67^. Here, in our modelling framework, we consider species to be facultative in the sense that they can survive and have positive growth rate in the absence of fitness benefits from their opposite guild of species.

Taking everything together, the population dynamics of animal species *i* can be written as

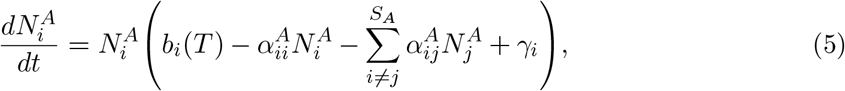

where *b*_*i*_(*T*)is the mean species growth rate for a given temperature *T*, 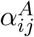, is the impact of competition by pollinator species *j* on species *i*, and *γ*_*i*_ is similarly the cumulative mutualistic impact of all plant species on pollinator *i*. Here, the exact formulation of 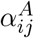 and *γ*_*i*_, and *b*_*i*_(*T*)is given in the supplementary section 1. Similarly, an analogous equation can be formulated for the plant species with index *A* replaced by *P*.

The evolutionary dynamics of the mean trait *µ*_*i*_ are given as:

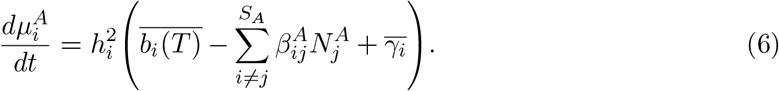

Here, 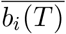 is the evolutionary impact of temperature on changes in the mean trait *µ*_*i*_, 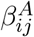 is the evolutionary impact of competition with species *j* on change in mean trait of species *i*, and 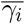 is the evolutionary impact of mutualistic trait-based interactions on changes in mean trait. Phenotypic variance in our modelling setup was given as 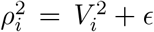, where *ϵ* is the environmental variance, and 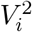 is the fixed genetic variance for the trait. Thus, the broad sense heritability, 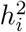 is fixed, with *ϵ* being fixed. Here, 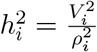 for a species *i* in question.

### 4.2 Numerical simulations and eco-evolutionary dynamics of plant-pollinator networks

Initial mean optimum phenotype values were drawn from a random uniform distribution of *U* [*T* +*b*_*w*_, *T* − *b*_*w*_], where *b*_*w*_ is the width of the temperature tolerance curve (see equation 4). This initial condition was chosen to ensure that species’ initial optimum values do not fall beyond the width of the tolerance curve given by *b*_*w*_. For details of parameter values used see table S1.

Initial species densities were set to 1, while the genetic variation, 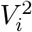, for each species was randomly drawn from a uniform distribution *U* [0.1, 0.25], and environmental variance for each species was randomly drawn from a uniform distribution *U* [0.009, 0.01]. We considered different plant-pollinator community sizes, consisting of 20, 30, and 40 species, with plants comprising 50% of the total species in each case. The initial network connectance and nestedness were determined by the plant-pollinator trait overlap (see section 3.1 below). However, the strength of such an interaction will be dependent on how close together their phenotypes are. We also assessed the strength of within-guild species competition and the impact of shape of the temperature tolerance curve. The strength of within-guild species competition was quantified by the width of the competition kernel given *w*_*c*_ in equation 3. We evaluated two levels of competition width *w*_*c*_: 0.05 and 0.15, with lower values indicating weak overall competition within guilds of a plant-pollinator network. We evaluated three different values of *a* of 0, 0.1, and 0.125.

In addition, with our modelling framework we also test the latitude-related diversity gradients, where we evaluate how plant diversity could potentially impact the architecture of plant-pollinator networks, in addition to evaluating the impact of temperature. We evaluated this mechanism by simulating communities with fixed pollinator richness while plant richness declined toward higher latitudes, and then quantifying network structure under different temperature regimes. Full details are provided in supplementary appendix S2.

### 4.3 Characterising plant-pollinator networks from simulated phenotype data

At *t* =0 and *t* =10^4^ time units, which was sufficient to get close to eco-evolutionary equilibrium (see fig. S17 for longer time units of simulations), we assessed key network properties, including connectance, nestedness, network specialization, and species rewiring, which are further described below. Our model is set up in a way that each pollinator species has a non-zero interaction rate with each plant species. However, if the mean optimum phenotypes are very far apart, the interaction rate will be small and not biologically meaningful. Thus, to characterise the emerging network structure, we need to quantify interaction strength and set a threshold for how strong the interaction needs to be to count as an edge in the network. We do this based on three different measures of interaction strength: the modified Gower similarity matrix weighted by phenotypic variance, which we call *Gower variance* ^68,69^, the Bhattacharya coefficient (*BC*) ^70,71^, and a true estimate (*TE*) based directly on the interaction rates in our model.

First, the modified Gower variance metric based on absolute mean trait difference between two species *i* and species *k* at any time *t* can be written as:

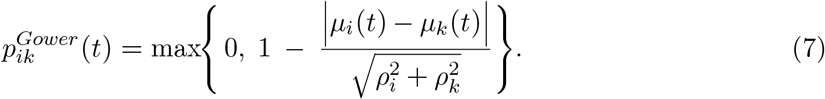

The classic Gower metric used in previous empirical studies^68,69^ does not account for phenotypic trait variances, but asks how similar mean trait values of species are in relation to the whole population sample. Our variance-normalised Gower metric in equation 7 divides by the interacting species’ trait variances and thus asks how similar the two species’ mean traits are relative to the widths of their trait distributions and then rescales the metric to lie between 0 (no overlap) and 1 (identical means). By taking the maximum, we ensure that the species whose mean-trait separation exceeds their joint standard deviation receive a similarity of zero. Ecologically, this means that species whose trait differences are small relative to their within-species variability are considered to share a high proportion of trait space, whereas those separated by more than their combined variance are treated as having no functional overlap.

Second, the Bhattacharyya coefficient quantifies the overlap between the Gaussian trait distributions of two species as ^70,71^:

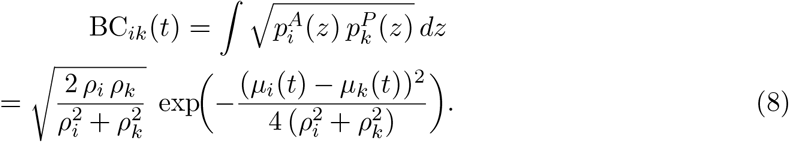

Ecologically, BC_*ik*_ then represents the proportion of shared optimum trait space i.e., the likelihood that their phenotypic optimum overlap sufficiently to allow meaningful interaction. The resulting similarity matrices, created from the above two similarity indices, captured the phenotype-based interaction strength between plant and pollinator species but did not take into account the mutualistic overlap *w* parameter.

Finally, we also create another trait-similarity metric directly from our model formulation which we call the true estimate *TE*, i.e.,

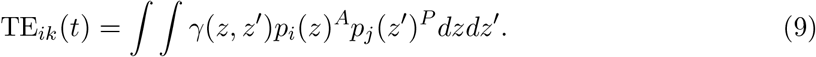

The three metrics are similar, but differ in some respects. The Gower variance and the Bhattacharya coefficient were solely based on the phenotype-matching hypothesis ^19,40^, however, *TE* takes the interaction kernel *γ*_*z,z*′_ into account as well. The hypothesis posits that the likelihood of species interaction is determined by the similarity of species mean traits, and their overlaps. The Gower variance similarity matrix is non-parametric, while the BC and TE similarity coefficient assumes that species trait distributions are Gaussian, which in our model simulations they are. Here, in the main text we show results network metrics calculated from the *Gower variance* because it is the one most commonly used in the literature^68,69^, but results of *BC*, and *TE* are in supplementary appendix.

Next, we created a binary interaction matrix from the pairwise interaction strength data and population abundances. Specifically, for the interaction matrix entry for species *i* and *k* to be 1, two conditions had to be met. First, both species’ densities had to be above a threshold of 10^−4^. Below this threshold we regard species as absent and any interaction involving them is essentially zero. This formulation ensures that even when a plant and a pollinator species have similar mean phenotypic traits, their actual interaction probability is influenced by whether they are present or effectively absent. Second, the interaction strength as quantified by equation 7, 8 or 9 had to be above an interaction-probability threshold. This threshold was fixed at 0.2 following previous empirical studies^68,69,72^ (but see robustness to two other different threshold values of 0.1, and 0.4 in appendix 3 Fig. S15-S16).

We then quantified network structure by computing connectance, nestedness based on the NODF metric, and overall network specialization (*H*2^*′*^) on the binary adjacency matrices using the `bipartite` R package ^73,74^. In parallel, we quantified phenotype-based rewiring over time by comparing interaction matrices between successive time points using Whittaker’s turnover index ^11^.

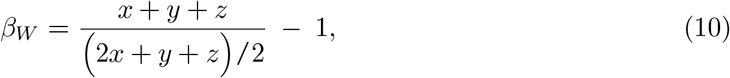

where *x* is the number of interactions shared between networks in two successive timepoints, and *y* and *z* are the interactions unique to each. Here, *β*_*W*_ can be further decomposed to *β*_*W*_ =*β*_*OS*_ +*β*_*ST*_, where *β*_*OS*_ is the rewiring part of interaction turnover, and *β*_*ST*_ is the compositional turnover. Here, we only were interested in rewiring aspect of interaction turnover and thus report that in our study, i.e., *β*_*OS*_. *β*_*OS*_ theoretically can range from 0 (no change) to 1 (complete rewiring) with higher values indicating greater rewiring in who interacts with whom. Next, we quantified median rewiring over time for each plant-pollinator network.

### 4.4 Plant-pollinator interaction data, and temperature data

We compiled species interaction matrices for 165 plant-pollinator networks: 91 networks from the Web of Life (www.web-of-life.es) interaction database across different bioregions of earth and the remaining 74 networks from open-access data ^10^. For each network location, we extracted latitude and longitude. Using these coordinates, we extracted climate variables from the WorldClim Global Climate Data version 2.1 at a spatial resolution of 30 arcseconds (~1 km), which captures fine-scale variation in temperature and precipitation ^75^. Specifically, we focused on mean annual temperature (MAT) as a key bio-climatic variable known to structure species distributions and interaction patterns across global gradients. Climatic data were extracted using the ‘terra’ package in R (v1.7-65) ^76^, ensuring consistent resolution and projection across all sites. These data are climatological averages from 1970–2001 which means they are long-term averages calculated from December 1970 through December 2001. From this temperature data, we specifically compiled MAT for each plant-pollinator network location. MAT ranged from as low as −5°C to as high as 30° C.

### 4.5 Statistical analysis of plant–pollinator network structure and temperature gradients

To reduce multicollinearity among plant–pollinator network descriptors, we conducted a Principal Component Analysis (PCA) on four commonly used metrics: modularity, connectance, nestedness, and network size. These metrics are often highly correlated due to underlying ecological constraints^56^. The PCA was performed on standardized (centered and scaled) values, and the first principal component (PC1), which explained the majority of variance (73.2%), was used as an integrated measure of overall network structure across sites. PC1 was positively correlated with nestedness and connectance (Pearson correlation coefficient 0.91 and 0.93, respectively) and negatively correlated with network size and modularity (Pearson correlation coefficient −0.78, and −0.85, respectively). PC2 explained a total of 18.1% variance (see figure S2).

We did a simple ordinary least squares (OLS) regression between PC1 scores and MAT as well as between network specialisation H2’ and MAT. In addition, we did an additional analysis where we do not use PCA axis scores but do an OLS regression of network metrics (connectance, modularity, nestedness, network-specialisation) against MAT (see figure S14). All analyses were conducted in R version 4.3.0^77^.

To account for spatial structure that might be present in our network data due to spatial correlation among network locations, we perform a spatial eigenvector mapping using R packages *spdep* and *spatialreg* ^36,59,78^ (see supplementary section 3 and figure S17).

## Acknowledgments

The authors would like to thank György Barabás for his comments on the manuscript. GB would like to acknowledge DFG Walter Benjamin grant no BA 7974/1-1 for funding this research. The authors declare no conflict of interest.

