## appendix for "Adaptive rewiring and temperature tolerance shape the architecture of plant-pollinator networks globally"

### 1 Mutualistic eco-evolutionary model

#### 1.1 Quantitative Genetics and Lotka-Volterra dynamics

We model the joint dynamics of pollinators and plants along a continuous phenotypic axis $z$. Each individual in the pollinator or plant guild is described by its trait value $z$, which varies across members of species $i$. Let $N_i^A(t)$ and $N_i^P(t)$ denote the abundances of pollinator and plant species $i$ at time $t$, respectively. The within species trait distribution is given by $p_i^{(A,P)}(z,t)$ with

$$\int_{-\infty}^{\infty} p_i^{(A,P)}(z,t)\,dz = 1. \tag{S1}$$

Under the quantitative-genetics assumption of many loci with small additive effects (weak selection)[1], trait values are normally distributed:

$$p_i^{(A,P)}(z,t) = \frac{1}{\sqrt{2\pi(\rho_i^{(A,P)})^2}} \exp\Big(-\frac{(z-u_i^{(A,P)}(t))^2}{2(\rho_i^{(A,P)})^2}\Big),$$

where $\mu_i^{(A,P)}(t)$ is the species mean trait and $(\rho_i^{(A,P)})^2$ its phenotypic variance (genetic + environmental).

Population and trait dynamics follow modified Lotka-Volterra equations with trait-based competition and mutualism. Adopting the framework from our own previous work and others[2–4], the per-capita growth rate of a pollinator individual with phenotype $z$ is

$$\begin{aligned} r_i^A(z) = {} & \underbrace{b(z,T)}_{\substack{\text{intrinsic}\\ \text{growth}}} - \underbrace{\alpha_{ii}^A N_i^A}_{\substack{\text{intra-}\\ \text{specific}\\ \text{competition}}} - \underbrace{\int \sum_{j=1;j\neq i}^{S_A} N_j^A\, \alpha(z,z''')\, p_j^A(z''',t)\, dz'''}_{\substack{\text{trait-based}\\ \text{inter-specific}\\ \text{competition}}} \\ & + \underbrace{\int \frac{\sum_{k=1}^{S_P} \gamma(z,z')\, A_{ik}\, N_k^P(t)\, p_k^P(z',t)}{1 + H \int \sum_{k=1}^{S_P} \gamma(z,z'')\, A_{ik}\, N_k^P(t)\, p_k^P(z'',t)\, dz''}\, dz'}_{\substack{\text{trait-based mutualistic}\\ \text{benefit (Type II)}}}\ . \end{aligned} \tag{S2}$$

An analogous expression holds for plant species, as:

$$\begin{aligned} r_i^P(z) = {} & \underbrace{b(z,T)}_{\substack{\text{intrinsic}\\ \text{growth}}} - \underbrace{\alpha_{ii}^P N_i^P}_{\substack{\text{intra-}\\ \text{specific}\\ \text{competition}}} - \underbrace{\int \sum_{j=1;j\neq i}^{S_P} N_j^P\, \alpha(z,z''')\, p_j^P(z''',t)\, dz'''}_{\substack{\text{trait-based}\\ \text{inter-specific}\\ \text{competition}}} \\ & + \underbrace{\int \frac{\sum_{k=1}^{S_A} \gamma(z,z')\, A_{ik}\, N_k^A(t)\, p_k^A(z',t)}{1 + H \int \sum_{k=1}^{S_A} \gamma(z,z'')\, A_{ik}\, N_k^A(t)\, p_k^A(z'',t)\, dz''}\, dz'}_{\substack{\text{trait-based mutualistic}\\ \text{benefit (Type II)}}}\ . \end{aligned} \tag{S3}$$

$b(z,T)$, is the per-capita growth rate of a phenotype $z$ as a function of temperature, $\alpha(z,z''')$ is the trait-dependent within-guild inter-specific competition kernel, and $\gamma(z,z')$ is the inter-guild Gaussian mutualistic kernel which designates that similar plant-pollinator phenotypes (z,z′ respectively) will have higher mutualistic benefits than others, $\alpha_i i$ is the intraspecific competition coefficient or density-dependence term. This coefficient is constant and fixed and phenomenological in the sense that this term captures all the unmodelled bits that could potentially impact population densities. Inclusion of this term also indicates that this is trait-independent. For instance, diseases, or unmodelled predation or herbivore on pollinators or plants which could potential restrict unbounded growth. H is the handling time.

### Population and trait dynamics

Integrating equation (S2) and equation (S3) over all trait values yields the coupled dynamics of population size and mean trait for species $i$[2,4,5]:

$$\frac{dN_i^A}{dt} = N_i^A(t)\int_{-\infty}^{\infty} r_i^A(z)\,p_i^A(z,t)\,dz. \tag{S4}$$

Substituting equation S2 into S4, we get

$$\begin{aligned}\frac{dN_i^A}{dt} = N_i^A(t)\int\Bigg(b(z,T) - \alpha_{ii}^A N_i^A - \int\sum_{j=1,i\neq j}^{S_A} N_j^A\alpha(z,z''')p_j^A(z''',t)dz''' + \\ \int\frac{\sum_{k=1}^{S_P}\gamma(z,z')A_{ik}N_k^P(t)}{1+H\int\sum_{k=1}^{S_P}\gamma(z,z'')A_{ik}N_k^P(t)p_k^P(z'',t)dz''}p_k^P(z',t)dz'\Bigg)p_i^A(z,t)dz\end{aligned} \tag{S5}$$

We can further write equation S5 as:

$$\frac{dN_i^A}{dt} = N_i^A(t)\left(b_i^A(T) - \alpha_{ii}^A N_i - \sum_{j\neq i}\alpha_{ij}^A N_j^A + \gamma_i\right) \tag{S6}$$

with temperature-dependent growth rate

$$b_i^A(T) = \int b(z,T)p_i^A(z,t)dz = \frac{g_i}{b_w - a\mu_i}\exp\Big(-\frac{(T-\mu_i)^2}{2(b_w-a\mu_i)^2+2\rho_i^2}\Big)\frac{b_w - a\mu_i}{\sqrt{(b_w-a\mu_i)^2+\rho_i^2}} - k_i, \tag{S7}$$

interspecific competition coefficient

$$\alpha_{ij}^A = \int\int\alpha(z,z''')p_j^A(z''',t)p_i^A(z,t)dzdz''' = \frac{\omega_c}{\sqrt{2\rho_i^2+2\rho_j^2+\omega_c^2}}\exp\Big(\frac{-(\mu_i-\mu_j)^2}{2\rho_i^2+2\rho_j^2+\omega_c^2}\Big)$$

and cumulative mutualistic effect

$$\gamma_i = \int\int\frac{\sum_{k=1}^{S_P}\gamma(z,z')A_{ik}N_k^P(t)}{1+H\int\sum_{k=1}^{S_P}\gamma(z,z'')A_{ik}N_k^P(t)p_k^P(z'',t)dz''}p_k^P(z',t)dz'p_i^A(z,t)dz, \tag{S8}$$

which we solved numerically.

Assuming the phenotypic trait is based on the quantitative genetic limit – i.e., it is controlled by many loci each of small additive effects under weak selection – the dynamics of the mean phenotype $u_i^{(A)}(t)$ of interest, here for the pollinators, can then be written as[3,4]

$$\begin{aligned}\frac{d\mu_i^A}{dt} = h_i^2\int(z-\mu_i^A)\Bigg(b(z,T) - \alpha_{ii}^A N_i - \int\sum_{j=1,i\neq j}^{S_A} N_j^A\alpha(z,z''')p_j^A(z''',t)dz''' + \\ \int\frac{\sum_{k=1}^{S_P}\gamma(z,z')A_{ik}N_k^P(t)}{1+H\int\sum_{k=1}^{S_P}\gamma(z,z'')A_{ik}N_k^P(t)p_k^P(z'',t)dz''}p_k^P(z',t)dz'\Bigg)p_i^A(z,t)dz,\end{aligned} \tag{S9}$$

where $h_i^2$ is the broad sense heritability of the trait. We can write this more compactly as

$$\frac{d\mu_i^A}{dt} = h_i^2\left(\overline{b_i(T)} - \sum_{j=1,i\neq j}^{S_A}\beta_{ij}^A N_j^A + \overline{\gamma_i}\right), \tag{S10}$$

where

$$\overline{b_i(T)} = \int(z-\mu_i^A)b(z,T)p_i^A(z,t)dz = \frac{g_i}{b_w - a\mu_i}\exp\Big(-\frac{(T-\mu_i)^2}{2(b_w-a\mu_i)^2+2\rho_i^2}\Big)\frac{\rho_i^2(b_w-a\mu_i)(T-\mu_i)}{((b_w-a\mu_i)^2+\rho_i^2)^{1.5}} \tag{S11}$$

represents the impact of direct selection on temperature tolerance and

$$\beta_{ij}^A = \frac{\rho_i \omega_c (\mu_j - \mu_i)}{(2\rho_i^2 + 2\rho_j^2 + \omega_c^2)^{1.5}} \exp(-\frac{(\mu_i - \mu_j)^2}{(2\rho_i^2 + 2\rho_j^2 + \omega_c^2)}) \tag{S12}$$

represents the impact of competition with other species in the guild on mean trait evolution. And finally

$$\overline{\gamma_i} = \int \int (z - \mu_i^A) \frac{\sum_{k=1}^{S_P} \gamma(z,z') A_{ik} N_k^P(t)}{1 + H \int \sum_{k=1}^{S_P} \gamma(z,z'') A_{ik} N_k^P(t) p_k^P(z'',t) dz''} p_k^P(z',t) dz' \Bigg) p_i^A(z,t) dz \tag{S13}$$

is the impact of mutualistic interactions on mean trait evolution, which is again evaluated numerically. Thus equation S5 and equation S10 represent population dynamics and the dynamics of the trait mean, respectively.

### 2 Plant diversity impact on plant-pollinator network architecture:

Our modelling results suggested that as temperature increased network specialisation decreased and network connectance increased, while controlling for total number of species, when $a > 0$, i.e., when there was a trade-off between species optimum trait and the width of their growth rate curve. This trade-off is empirically observed in species across a latitudinal gradient[(6)]. We corroborated our modelling results with empirical plant-pollinator networks from across the globe. Our empirical analyses reveal that as mean annual temperature (MAT) rises, mutualistic networks become less specialized and more densely interconnected and nested.

Earlier work has linked this latitudinal gradient in specialization to elevated plant richness in tropical regions where abundant floral resources dilute per-species encounters and favor broad-diet (generalist) foraging[(7)]. In contrast, cooler, temperate communities with sparser floral assemblages tend to constrain pollinators to narrow interaction niches, reinforcing higher specialization. We tested this particular hypothesis in our model simulations where we fixed the total number of pollinators to just 10 species and varied plant richness across a temperature gradient mimicking the scenario where in temperate regions there are less plant species and in tropical latitudes there are more plant species. This relationship was linear and was modelled to be $y = 4 + 0.5x$, where $y$ is the plant richness, and $x$ is the temperature. This specific scenario was to test the hypothesis of latitudinal plant-diversity gradient driving network generalisation in the tropics[(7)]. The temperature gradient went from 0° C to 30° C, and thus plant richness varied as temperature increased, such that when temperature was 0° C, the number of plant species was 4 and when temperature was 30° C, plant richness was 19 (rounded to whole number). If plant diversity per pollinator species was the main driver for generalisation observed in low latitudes or higher temperatures, we should observe a negative relationship between temperature and specialisation even when $a = 0$, i.e., there is no trade-off between species peak tolerance and width of the tolerance curve.

In figure S4, we observe the opposite for $a = 0$. As temperature increased, networks overall became more specialised rather than becoming generalised indicating that as plant richness increased, networks became more specialised, and connectance decreased despite different strength in species competition (Figure S4B, $a = 0$). However, when $a > 0$, we observe that despite increases in plant richness, we still observe negative relationship between H2′ and temperature indicating that species peak tolerance and width trade-off, competition, was a strong driver of network generalisation in high temperatures.

### 3 Spatial detrended analysis of network structure and MAT:

To examine the relationship between network structure and climatic conditions while accounting for potential spatial dependence among sites, we conducted spatial eigenvector mapping analyses using the *spdep* and *spatialreg* packages in R. We first compiled a global dataset of 165 plantpollinator networks with associated climatic variables, including mean annual temperature (MAT). For each network we calculated standard descriptors (connectance, nestedness, modularity, and network size), and performed a principal component analysis (PCA) on these metrics. The first principal component (PC1) captured the main variation in network architecture, and was used as the focal response variable in subsequent regressions. Pearson correlations confirmed that PC1 was strongly associated with each individual network property.

Table S1: Description of parameters and their values used in the study. $U[a, b]$ is the uniform distribution between $a$ and $b$.

| Symbol | Description |
|---|---|
| $V_i^2$ | Genetic variance of species $i$. |
| $\rho_i^2$ | Phenotypic variance of species $i$. |
| $k_i$ | intrinsic mortality of species $i$, fixed at 0.1. |
| $a$ | controls the trade-off between peak tolerance and width of the thermal tolerance, various from 0, 0.1, 0.125. |
| $h_i^2$ | Heritability of species $i$'s trait; equal to $\frac{V_i}{\rho_i}$. |
| $g_i$ | modulates the peak of temperature-dependent growth rate; fixed at 1.5 for all species. |
| $\alpha_{ii}$ | trait-independent intraspecific competition; fixed at 1 for all species. |
| $b_w$ | modulates the trade-off between temperature-dependent growth rate width, and peak; fixed at 4.5 for all species. |
| $\alpha_{ij}^{(A,P)}$ | pairwise inter- and intraspecific competition; intraspecific competition $\alpha_{ii}$ was fixed at 1 for all species, whereas interspecific competition $\alpha_{ij}$ was derived from equation **??** (see appendix 1). |
| $\gamma_0$ | maximum mutualistic interaction strength; fixed at 1. |
| $w$ | width of mutualistic Gaussian interaction kernel; fixed at 1. |
| $w_c$ | width of the Gaussian competition kernel, either 0.05, or 0.15. |
| $k_i$ | mortality from the temperature tolerance curve; fixed at 0.1 for all species. |
| $T$ | Average local environmental temperature, which varied from $0°C$ to $29°C$. |
| $u_i$ | Mean optimum trait values of species. |
| $N_i^A$, $N_i^P$ | Population density of animal and plant species. Their initial value was 1 for all species. |
| $\epsilon$ | environmental variance, randomly sampled for all species from $U[0.009, 0.01]$. |
| $H$ | Handling time, fixed at 0.25 for all species. |

Because spatial proximity of sites can generate autocorrelation in network properties, we constructed spatial weights matrices using geographic coordinates (latitude and longitude) of each network. Neighbors were defined with a distance band of 05000 km, and a row-standardized spatial weights list was generated. Morans I tests indicated significant spatial autocorrelation in the residuals of initial ordinary least squares (OLS) regressions of PC1 against MAT ($p < 0.05$). We therefore applied the Moran eigenvector spatial filtering approach (*Spatial Filtering* function), which selects a subset of orthogonal eigenvectors representing broad- to fine-scale spatial patterns. These eigenvectors were incorporated as additional predictors alongside MAT in OLS models, thereby removing spatial autocorrelation from the residuals. Next, after controlling for spatial structurem , we plotted spatially controlled MAT against PC1, see figure S17.

### 4 Supplementary Figures:

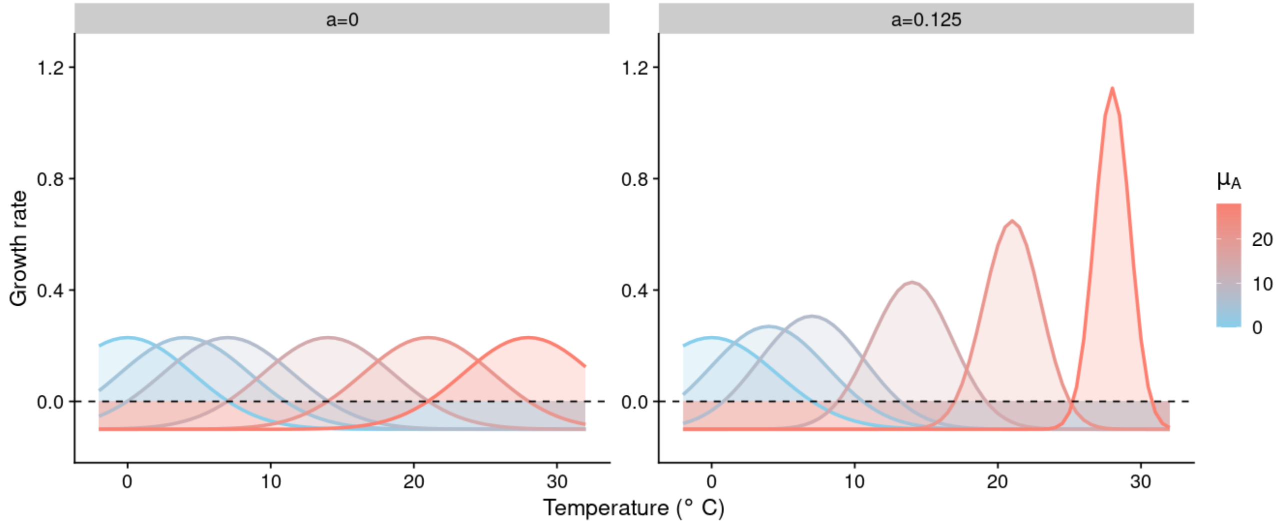


Figure S1: Growth rate or thermal tolerance of different species having different mean optimum trait going from as low as 2° to as high as 28°. When $a = 0$, there is no trade-off between peak of the thermal tolerance and width of the thermal tolerance curve, i.e., a species which occurs at low temperatures has similar growth curve as a species that occurs at higher temperatures. When $a = 0.125$, there is a trade-off between peak of the tolerance curve and width of the curve, indicating that species that occur at low temperatures do not have the same rate of growth compared to species that occur at higher temperatures. Parameters are $b_w = 4.5$, $g = 1.5$, $k_i = 0.1$, for equation 4 in the main-text.

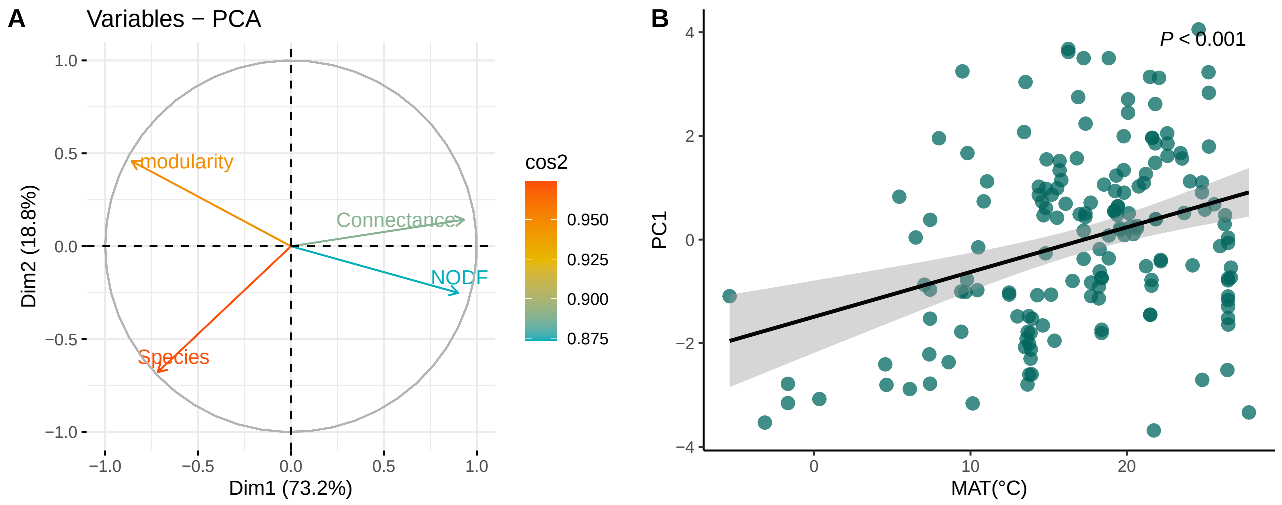


Figure S2: Principal component analysis of four network metrics: nestedness (NODF), number of species, connectance, and modularity of 165 empirical plant-pollinator networks collated. (A) PC1 is strongly correlated positively to connectance and nestedness and negatively to modularity. (Right) Positive relationship between PC1 and mean annual temperature (MAT). As MAT increases, PC1 increases indicating that nestedness (NODF), and connectance increases, but network size decreases.

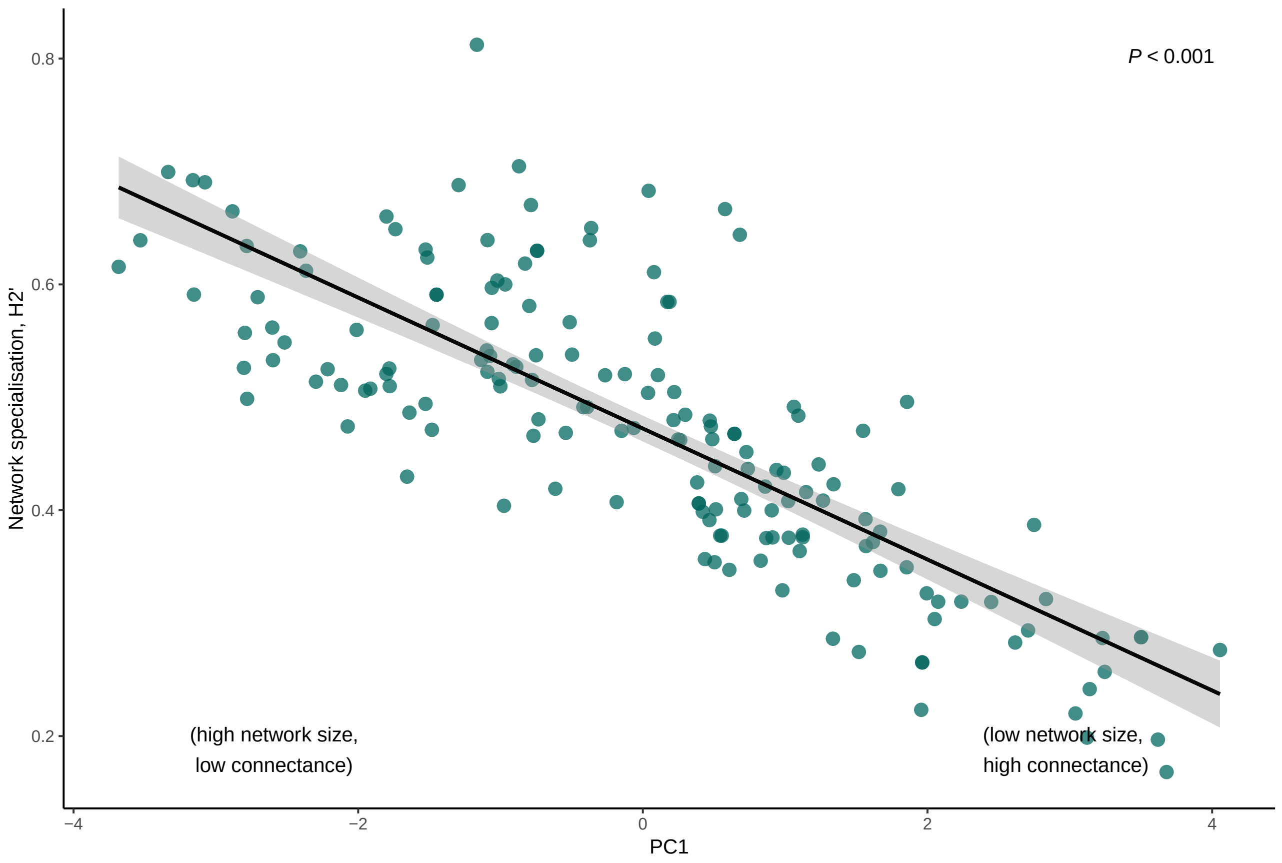


Figure S3: Relationship between network specialisation , H2′ and Principal component axis 1 scores, PC1. As PC1 increases (i.e., high connectance but low network size), network specialisation decreases, i.e., species become more general.

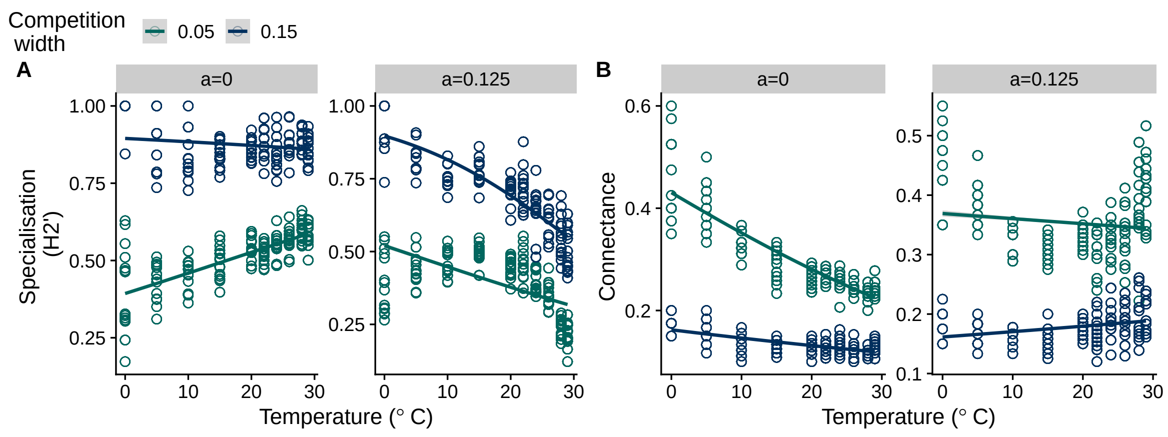


Figure S4: Network specialisation and connectance in relation to environmental temperature with variable plant richness in simulated plant-pollinator networks. Here, as temperature increases, plant richness increases linearly from 4 species at 0° C to 19 species at 30° C, but pollinator richness was capped at 10 species. A) Specialisation still decreased regardless when $a > 0$, but when $a = 0$, specialisation increased. B) Connectance decreased as temperature increased when $a = 0$, but when $a > 0$, connectance increased at high temperature as in the main text.

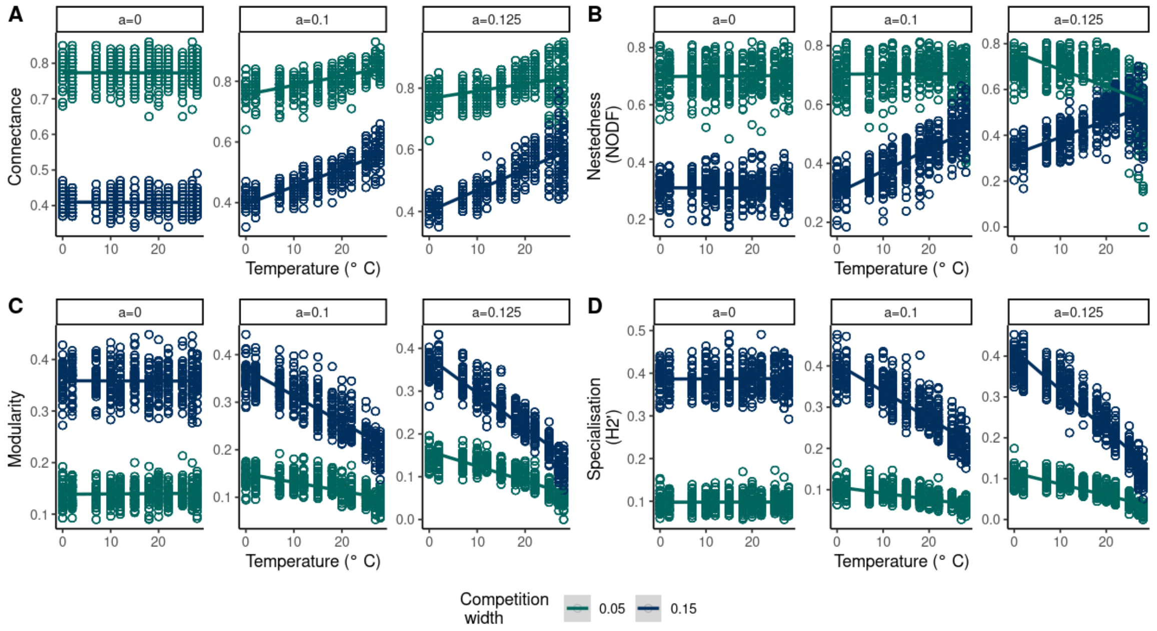


Figure S5: Adaptive phenotypic species rewiring and tolerance to temperature impacts network specialisation and structure of plant-pollinator networks of 20 species. A) As temperature increases, final equilibrium network connectance when $a > 0$, with strong competition ensuring an overall decrease in network connectance. However at $a = 0.125$, for lowest competition width, we observe nestedness to slightly fall off at high temperature values. C) Similarly, nestedness increases as temperature increases when $a > 0$, D) Similarity, modularity was negatively correlated with temperature when $a > 0$, and finally E) H2 i.e., network specialisation, decreases as temperature increases when $a > 0$, with competition having an impact more on the intercept and less on the slope of the relationship. Line's represent generalised linear model with 95 % confidence interval. Network metrics such as connectance, nestedness, modularity, and H2 are estimated using the BC metric. Starting network size is 20. Parameters as in Table S1.

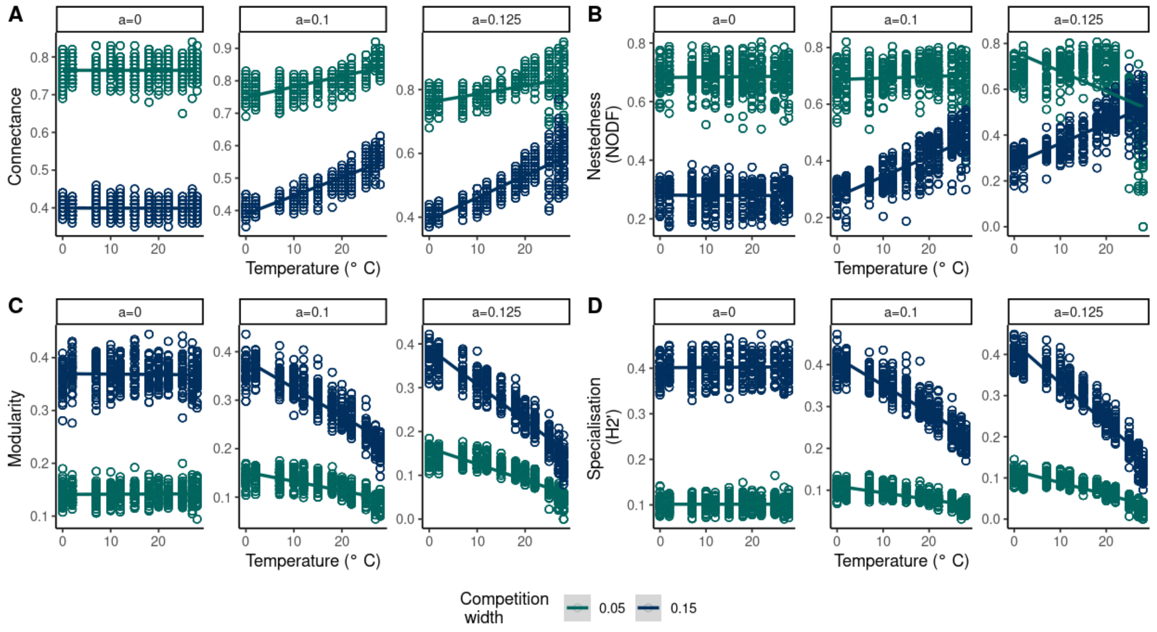


Figure S6: Same as figure S5 but the network adjacency matrix was quantified by the true estimate metric for a network size of 20 species. Parameters as in Table S1.

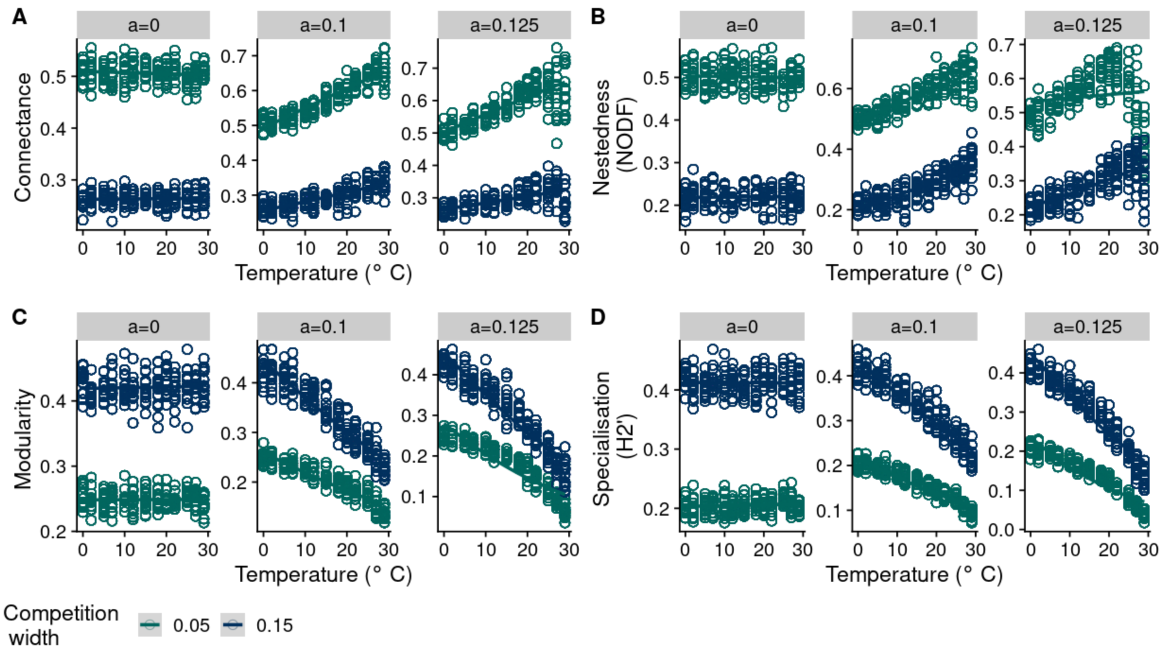


Figure S7: Same as figure S5-S6 but for plant-pollinator networks of 40 species, and network adjacency matrix was quantified by BC metric. Parameters as in Table S1.

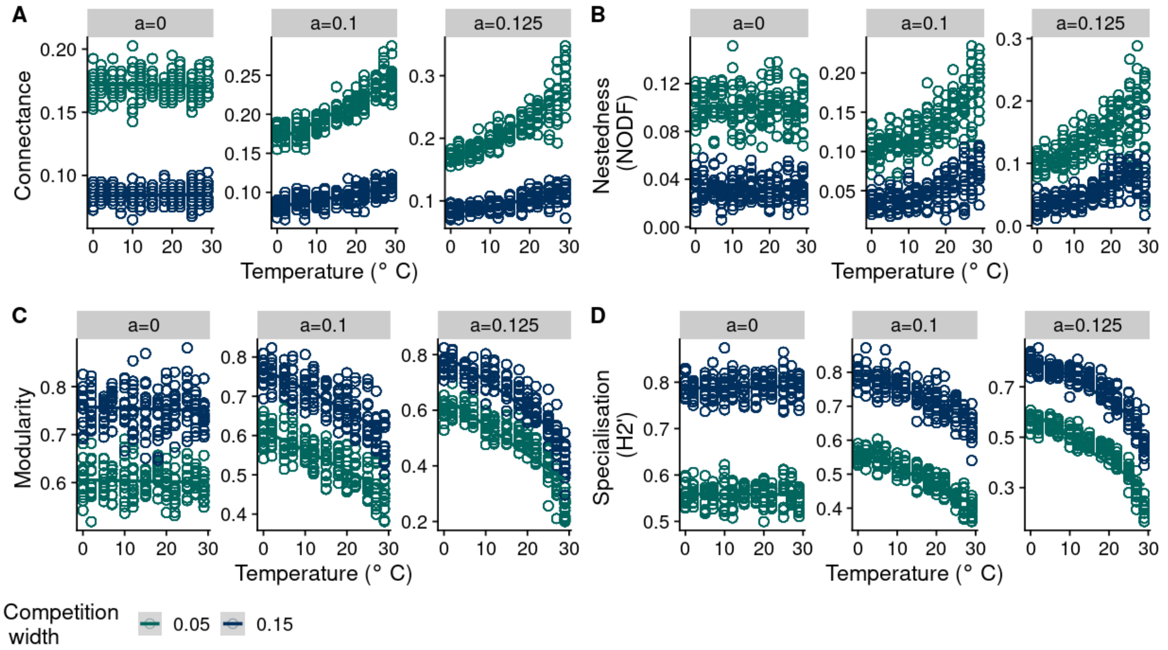


Figure S8: Same as figure S5-S6 but for plant-pollinator networks of 40 species, and network adjacency matrix was quantified by Gower variance metric. Parameters as in Table S1.

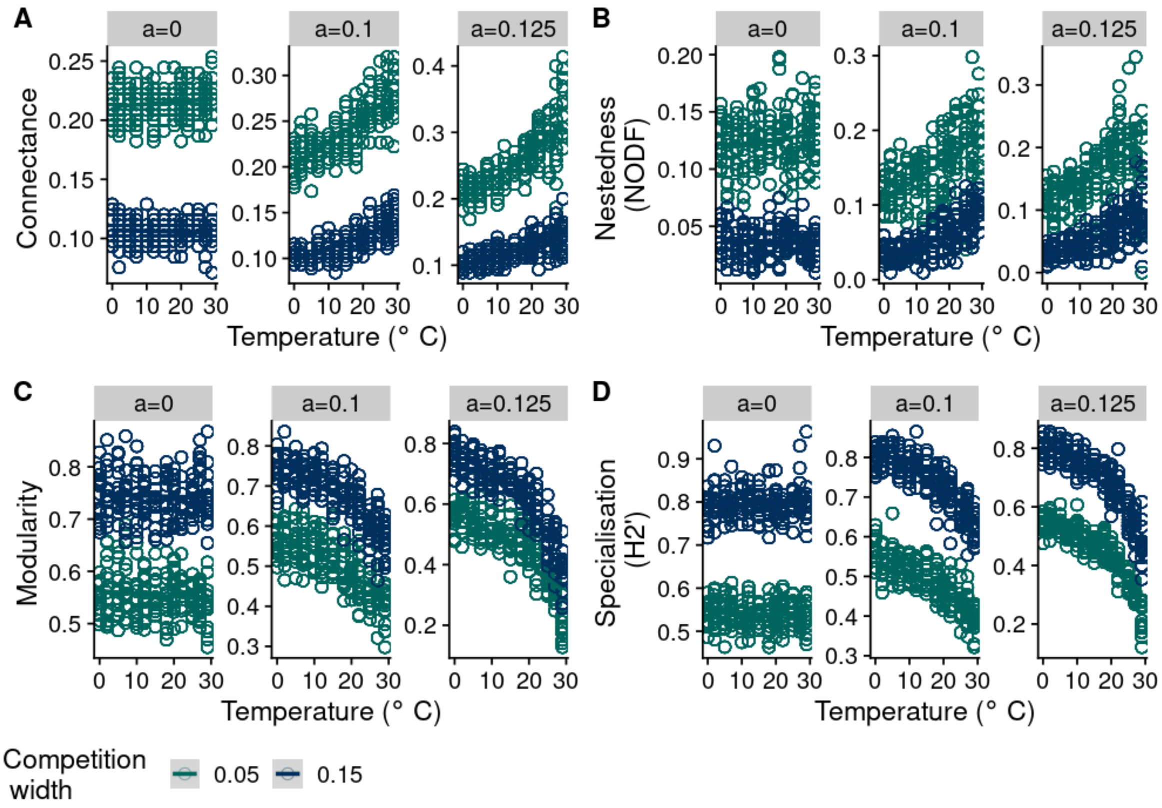


Figure S9: Same as in figure S5-S6 but for plant-pollinator network of size 30 species, and network adjacency matrix was quantified by Gower variance metric. Parameters as in Table S1.

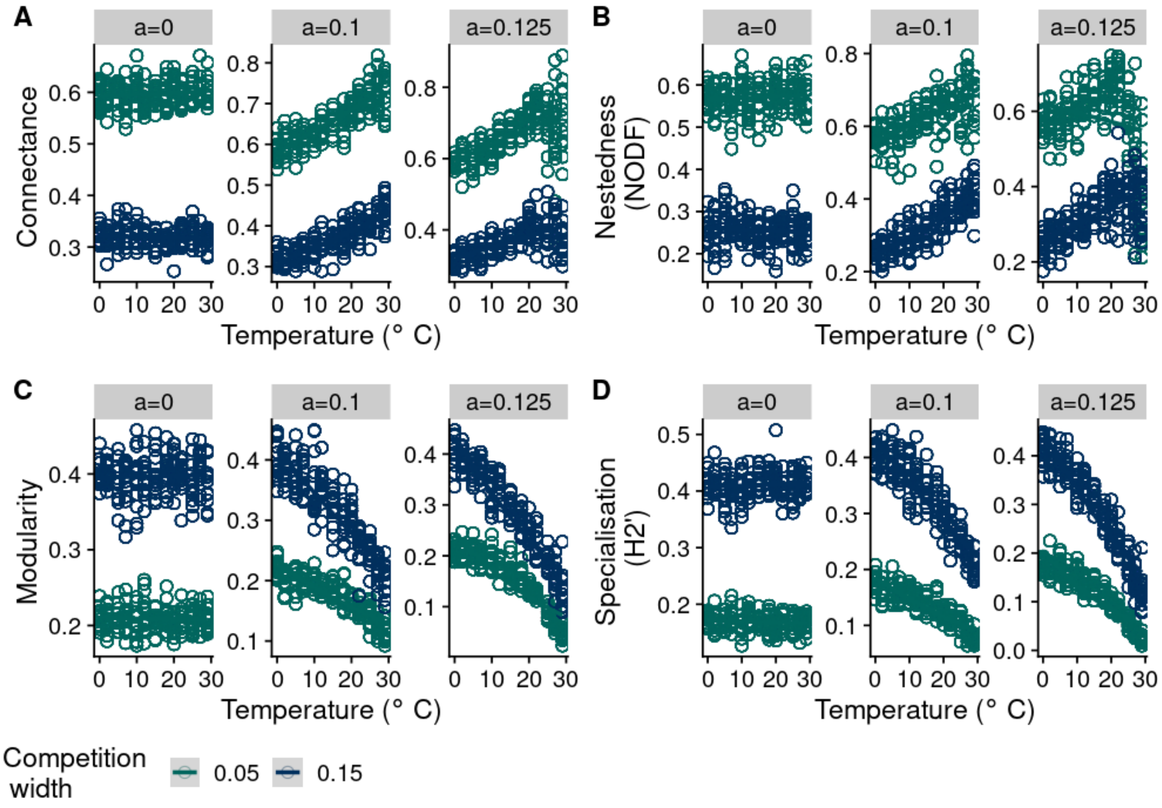


Figure S10: Same as in figure S5-S6 but for plant-pollinator network of size 30 species, and network adjacency matrix was quantified by BC metric. Parameters as in Table S1.

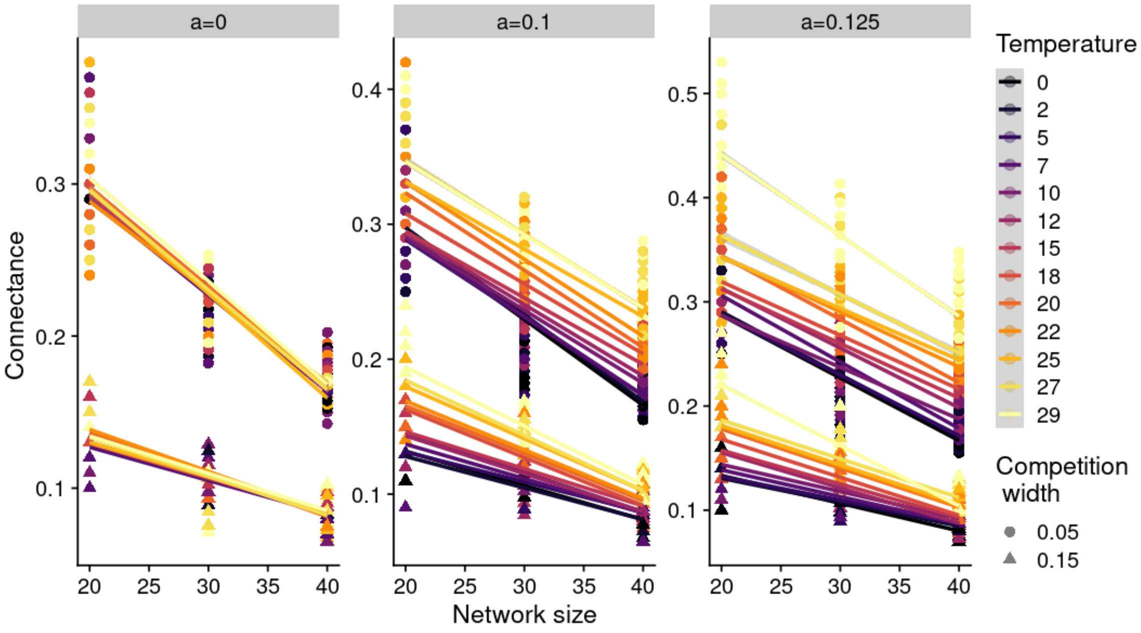


Figure S11: Relationship between starting network size and final network connectance in our model simulations for different temperature regimes and competition width, and tolerance curve, $a$. As in empirical networks observed globally, connectance has a negative relationship with network size in our simulations too across different levels of $a$.$a = 0$: there is no variability observed in network connectance and network size relationship. $a = 0.1$: we observe temperature to have an impact on the intercept values and slightly on the slope of the relationship. We observe that smaller networks at higher temperatures have more connectance than larger networks at higher temperatures. $a = 0.125$: we observe similar negative relationship between network size and connectance impacted by temperature. Strength of competition (given by triangle shape points) i.e., competition width $0.05, 0.15$ impacted final connectance, with higher competition having low connectance overall. Parameters as in Table S1.

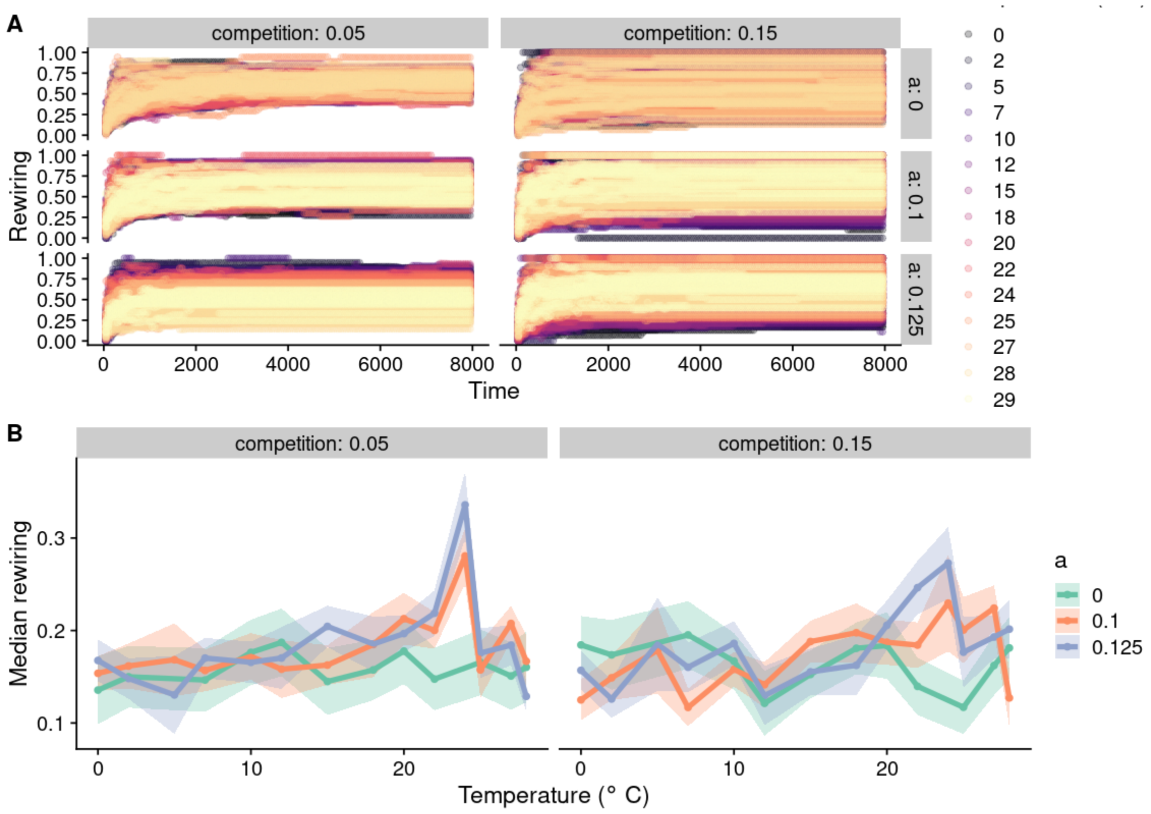


Figure S12: Dynamics of adaptive phenotypic species rewiring with different initial conditions compared to Fig. 3. Here, mean species trait values at $t = 0$ were sampled from a random uniform distribution $U[T - 2.5, T + 2.5]$, where $T$ is the temperature regime. In the main text, they were sampled from $U[T - 4.5, T + 4.5]$ as temperature width $b_w$ was fixed at 4.5. A) Rewiring over time increases for different temperatures and strength of competition. B) Median phenotypic based rewiring in relation to environmental temperature, competition, and shape of the temperature tolerance curve. Rewiring on average is slightly higher when $a > 0$. Lines represent median rewiring per environmental temperature and shaded areas represent 95% confidence interval. Starting connectance was 1 and network size was 20 species. Parameters as in Table S1.

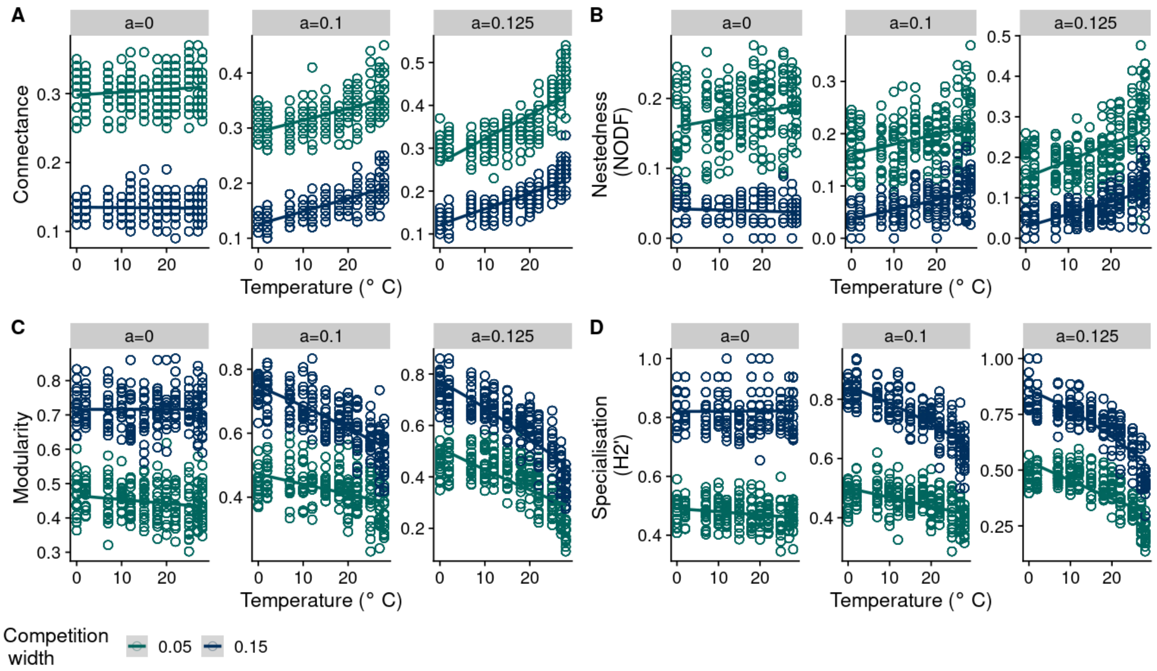


Figure S13: Same as figure S9 but initial starting mean trait values are sampled from $U[T - 2.5, T + 2.5]$ for 20 species plant-pollinator network. Parameters as in Table S1.

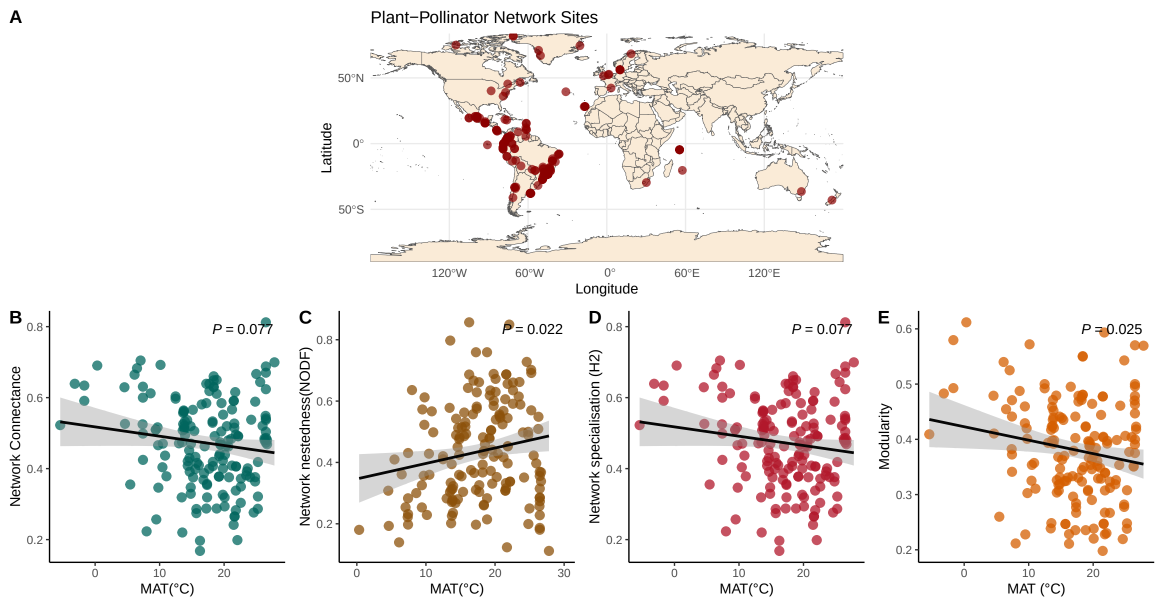


Figure S14: Network architecture of plant-pollinator networks in relation to mean annual temperature. A) Geographic location (latitude and longitude) of 165 plant-pollinator networks. (B) Relationship between network connetance and mean annual temperature (MAT), C) nestedness (NODF), D) network specialistion (H2′), E) modularity. Each point represents a plant-pollinator network. Lines indicate fitted linear regressions with 95% confidence intervals.

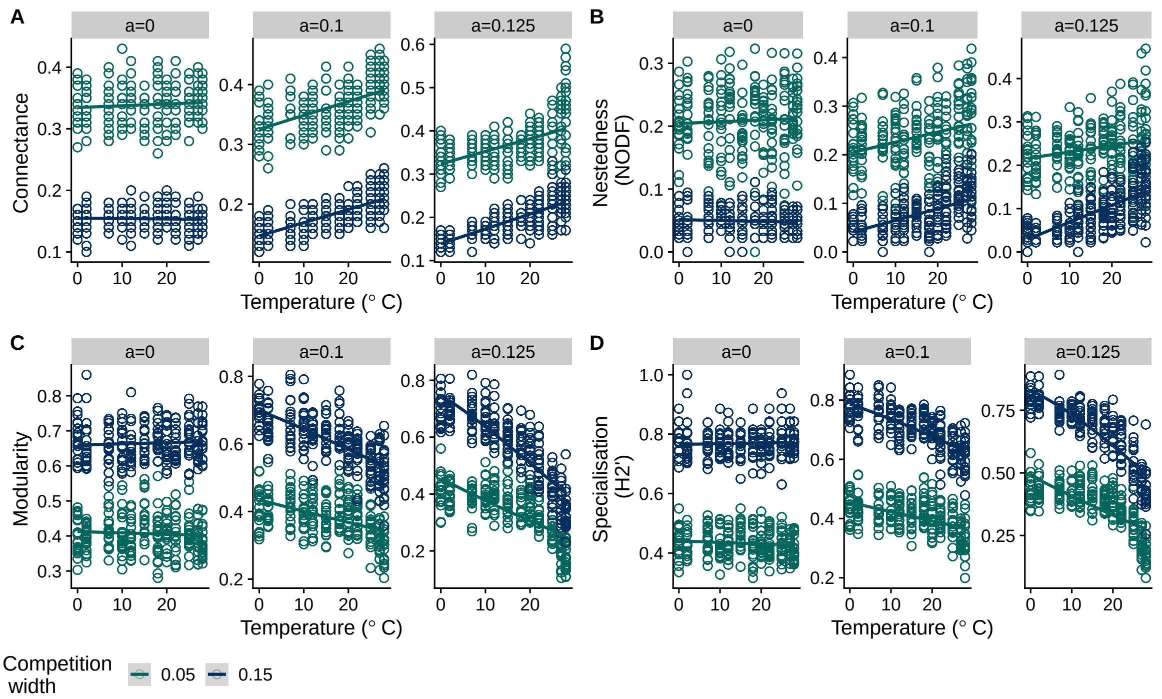


Figure S15: Same as figure S5 with plant-pollinator network size of 20 species, with Gower variance determining the adjacency interaction matrix and interaction threshold of 0.1. Parameters as in table S1.

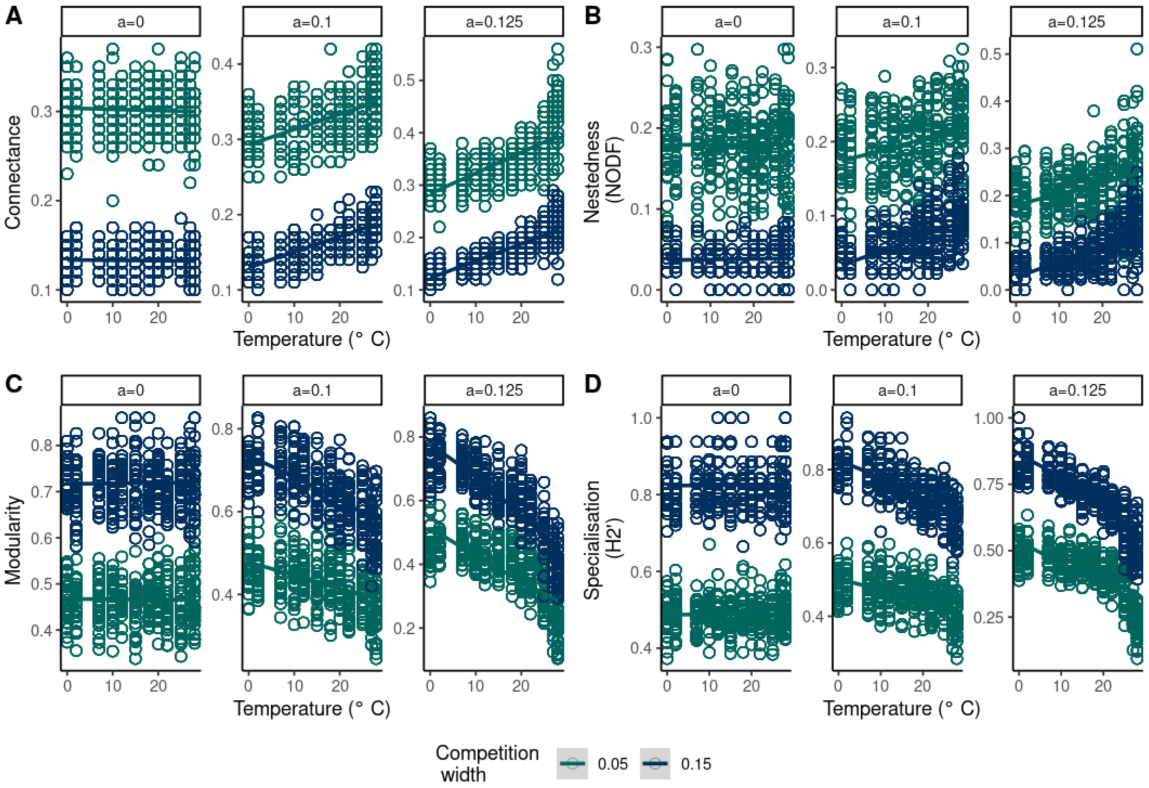


Figure S16: Same as figure S15 with plant-pollinator network size of 20 species, with Gower variance determining the adjacency interaction matrix and interaction threshold of 0.4. Parameters as in table S1.

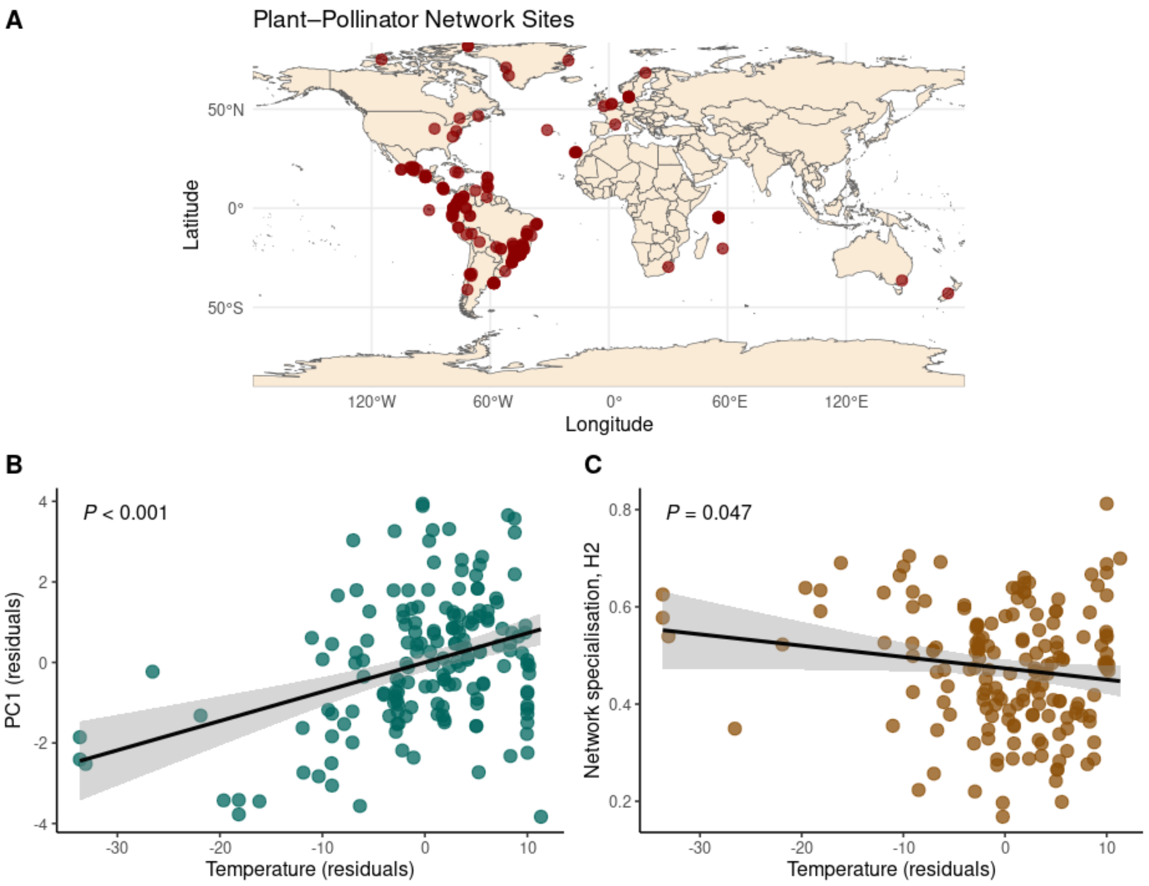


Figure S17: Same as figure S14, but after taking spatial correlation among network sites and then plotting the residuals of PC1 against residuals of MAT in B, and Network specialisation against residuals of MAT in C.

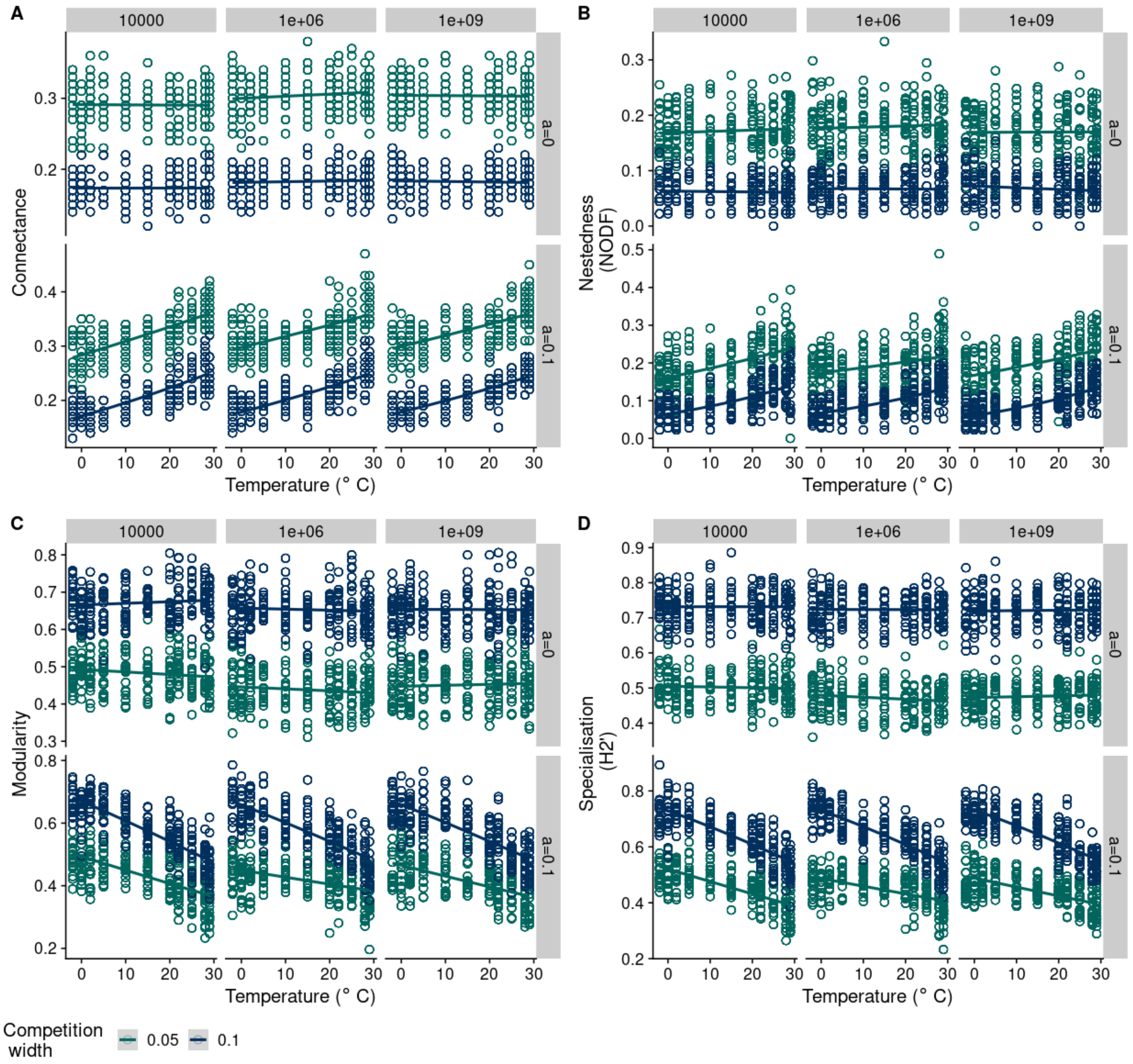


Figure S18: Same as figure S15, but for different time points of eco-evolutionary simulation i.e., total time points of simulations were $t = 10000$, $t = 10^6$, and $t = 10^9$, for two different $a$ values and plotted for network metrics of connectance, nestedness, modularity and network specialisation against temperature and for different competition width. Take away message from this plot is that total time for simulation had no significant impact on the outcome of the results.
